# Metastatic Osteosarcoma is Characterized by Loss of Osteoblastic Lineage Fidelity

**DOI:** 10.64898/2026.06.10.731352

**Authors:** Yutaro Tanaka, James J. Morrow, Salvador Casaní-Galdón, Erica M. Pimenta, Jiayi Li, Byron Butaney, Chadi A. El Farran, Julie Karam, Joshua P. D’Antonio, Kevin Bi, Laura Valderrábano, Sabrina Y. Camp, Yun Jee Kang, Shreya Johri, Lester Heredia Gopar, Shreya Raman, Breanna Titchen, Jingxin Fu, Josephine Yates, Samantha E. Hoffman, Theodora Pappa, Emma Su, Jihye Park, Amanda E. Garza, Anwesha Nag, Aurelia R. Reynolds, Mark Connelly, Sébastien Vigneau, Aaron R. Thorner, Megan E. Anderson, Jovana Pavisic, Brian D. Crompton, Eliezer M. Van Allen, Katherine A. Janeway, Natalie B. Collins, Bradley E. Bernstein, Riaz Gillani

## Abstract

Osteosarcoma (OS) is a rare and aggressive bone cancer with limited therapeutic progress in several decades. To characterize the transcriptional diversity of malignant cell states in OS, we analyzed single-nucleus RNA sequencing data from 24 tumors, including both primary and metastatic lesions. We identified a canonical osteoblastic cell state that predominated in primary tumor cells, but was dampened in metastases. Integrating tumor data with data from embryonic skeletal development and an experimental osteoblast differentiation model revealed that metastatic OS cells resemble poorly specified mesenchyme enriched for non-osseous programs. Clonal populations in metastatic samples acquired mesenchymal cell states distinct from paired primary tumors. Similar cell state shifts were evident in post-treatment primary tumors relative to paired pre-treatment samples. Our results suggest that localized post-treatment and metastatic OS malignant cells lose osteoblastic lineage fidelity, which may represent a unifying axis of disease evolution and reveal new therapeutic vulnerabilities.

## Introduction

Osteosarcoma (OS) is the most common bone malignancy, predominantly impacting children and adolescents. Therapeutic options and clinical outcomes for patients have remained largely unchanged for over four decades (1). Although advances have been made in nominating prognostic biomarkers, the most reliable predictor of patient outcome remains the presence of clinically detectable metastases at diagnosis (2; 3; 4). Patients who progress, relapse post-therapy, or develop metastatic disease face a starkly poorer prognosis, with a 5 year event-free survival of around 20%, compared to approximately 60% in those with localized, treatment-responsive disease (5; 6). Although various genetic, epigenetic, and signaling pathway alterations have been associated with disease progression in OS, the shared and distinct biology of localized and metastatic disease has not been thoroughly characterized (1; 7; 8).

A growing body of evidence across solid tumor types suggests that dysregulated cellular differentiation, aberrant developmental trajectories, and altered cellular states are pivotal drivers of tumorigenesis, therapeutic resistance, and metastasis (9; 10; 11; 12; 13). Specifically, prematurely arrested differentiation or active regression towards a less differentiated state (dedifferentiation), has been shown to enable malignant cells to bypass normal developmental constraints, sustain uncontrolled proliferation (14), and acquire the adaptive plasticity required to achieve metastatic progression (15). However, in OS, the developmental states that underlie tumor initiation and metastasis remain poorly characterized (16; 17). This may be due, in part, to an incomplete understanding of the transcriptional dynamics regulating the normal developmental trajectory of the bone-forming osteoblastic lineage.

While recent scRNA-seq based studies have begun to characterize primary OS tumor cell states in relation to transcriptional phenotypes seen in normal osteoblastic differentiation (18; 19; 20; 21; 22), metastatic disease remains largely underexplored due to data scarcity. To gain deeper insights into the differential biology of localized and metastatic OS, we analyzed single-nucleus RNA sequencing (snRNA-seq) data from OS tumor samples, including pre-and post-treatment specimens and a significant number of metastatic tumors. To evaluate the malignant OS cell states observed in the context of normal differentiation and development, we mapped the differentiation trajectory of primary human mesenchymal stem cells (MSCs) toward osteoblasts and leveraged a human embryonic skeletal development atlas. Through an integrative analytical approach, we characterized heterogeneous malignant cell populations in OS and uncovered a loss of osteoblastic lineage fidelity in post-treatment and metastatic OS.

## Results

### snRNA-seq reveals a loss of osteoblastic markers in metastatic OS tumors

We analyzed snRNA-seq of 24 OS tumor specimens from 18 pediatric patients (age range 4-21) **(Supplementary Table 1)**. This cohort included specimens from 7 metastatic and 17 primary tumors with two primary-metastatic tumor pairs and two primary pre-post treatment pairs **(Fig. 1a)**. All samples passed stringent quality controls and diagnoses were reconfirmed by pathology review **(Methods / Supplementary Table 2)**. We assigned cells (n=102,956) to broad cell type classes using a supervised non-negative matrix factorization approach **(Methods / Supplementary Material 1 / Extended Data Fig. 1a-b)**. We performed further granular cell type annotation for each broad tumor microenvironment cell type class, identifying 16,635 immune, 16,669 endothelial, and 10,100 epithelial cells **(Supplementary Material 2 / Supplementary Table 3)**.

**Fig. 1:**
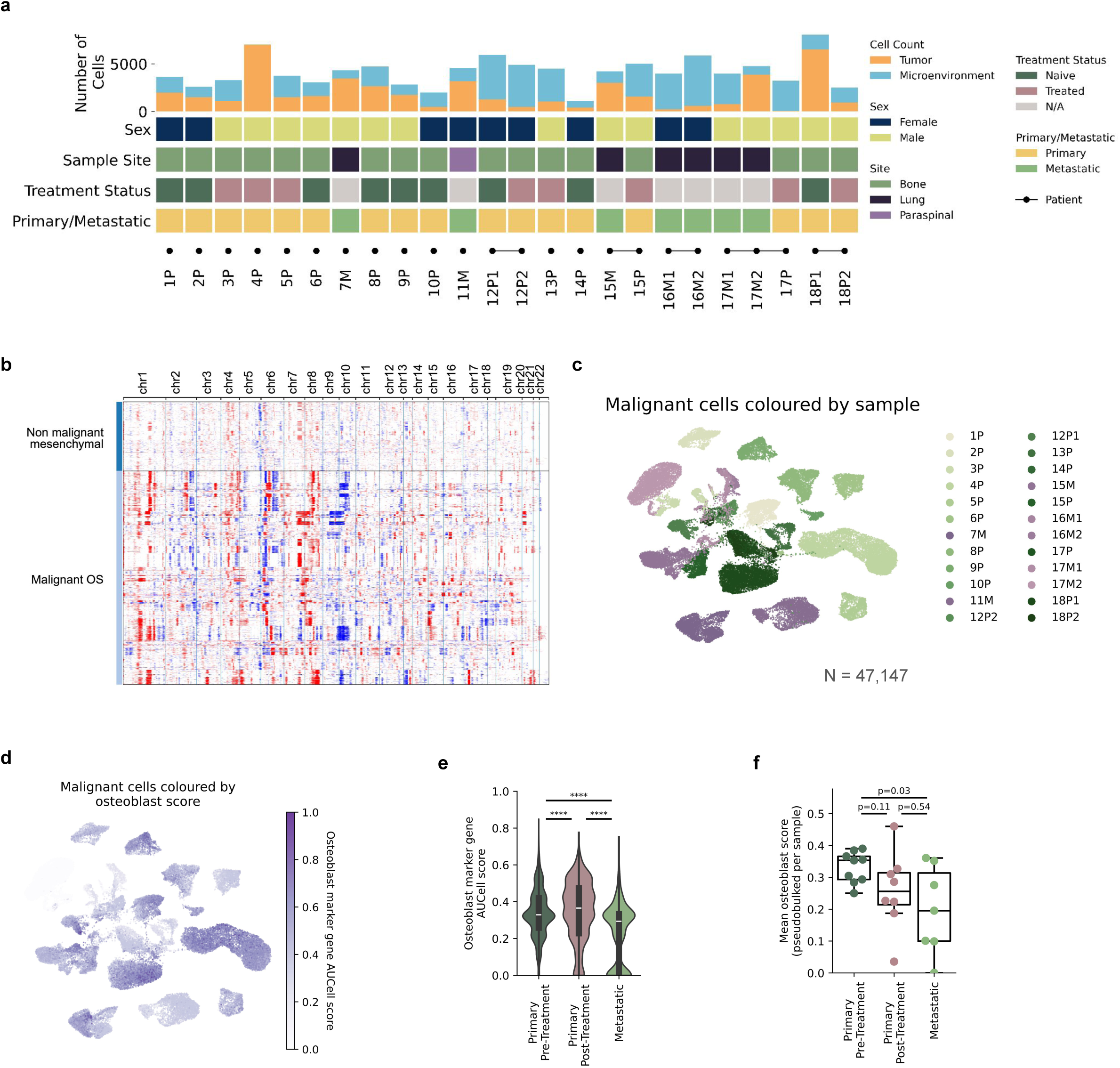
snRNA-seq reveals a loss of osteoblastic markers in metastatic OS tumors (a). Clinical metadata and cellular composition of the osteosarcoma (OS) cohort (n = 24 samples). The top stacked barplot details the counts of malignant versus microenvironment cells per sample, accompanied by annotation tracks summarizing patient sex, anatomical site, treatment status, and disease stage (primary/metastatic). **(b).** Inferred copy number alterations across autosomes of cells identified as non-malignant or malignant OS cells (n = 47,147). **(c).** Uniform Manifold Approximation and Projection (UMAP) of the tumor cells in the cohort, colored by sample. **(d).** UMAP of the tumor cells in the cohort, Osteoblast marker gene signature (*COL1A1 / COL1A2 / RUNX2 / MMP9 / IBSP / SPP1 / BGLAP / ALPL / SP7*) AUCell score. **(e).** Distribution of Osteoblast marker gene signature AUCell scores between primary pre-treatment, primary post-treatment, and metastatic tumor of origin, and **(f)** collapsed by sample mean. All statistical tests were performed as two-sided Mann-Whitney U tests (**** denotes p < 0.0001).

To identify malignant OS cells, we inferred copy number alterations across all autosomal chromosomes and filtered for mesenchymal cells with significant copy number alterations **(Fig. 1b)**, finding 47,147 OS cells across 24 samples. We found that the proportion of malignant and non-malignant cell types was highly variable across tumors in the cohort **(Extended Data Fig. 1c)**. Within the malignant cell population, we found that cells predominantly clustered by patient, indicating a high degree of inter-patient gene expression heterogeneity consistent with previous observations in OS **(Fig. 1c / Extended Data Fig. 1d)**(18). This observation of inter-tumor transcriptional heterogeneity was further confirmed by limited correlation of the top 3,000 highly variable genes across the cohort **(Extended Data Fig. 1e)**.

To investigate the transcriptional phenotypes present within malignant OS cell populations, we first evaluated an aggregate score of 9 literature-derived osteoblast marker genes across all malignant cells, observing a broad gradient of expression **(Fig. 1d)**. We found that metastatic tumor cells had lower aggregate osteoblast marker gene expression compared to primary tumor cells stratified by samples pre-and post-treatment **(Fig. 1e / Extended Data Fig. 1f)**. Complementary cell-rank enrichment analysis demonstrated primary tumor cells were enriched at the high-scoring extreme of the osteoblastic score gradient, whereas metastatic tumor cells were enriched at the low end of the spectrum **(Extended Data Fig. 1g)**. We also evaluated aggregate osteoblastic marker gene expression at the sample level, and observed a significantly lower expression in metastatic samples relative to naive primary samples **(Fig. 1f)**.

### Localized and metastatic OS malignant cells adopt heterogeneous mesenchymal gene expression programs

To further characterize the transcriptional cell states in localized and metastatic OS cells, we next adapted an iterative, unsupervised approach to define gene expression programs (GEPs)(23; 24; 25) **(Fig. 2a / Methods)**. We first derived GEPs in each patient tumor sample, removed any low usage GEPs, then clustered the remaining programs to identify and collapse highly similar GEPs into metaprograms (MPs). This approach identified 10 MPs that were present across tumor samples **(Fig. 2b / Supplementary Table 4 / Supplementary Material 3)**. In subsequent analyses, we characterized the activity of each MP using the MP’s 100 most active genes. We found that the majority of tumor cells expressed a small number of programs (mean 1.37±1.23) **(Extended Data Fig. 2a)**.

**Fig. 2:**
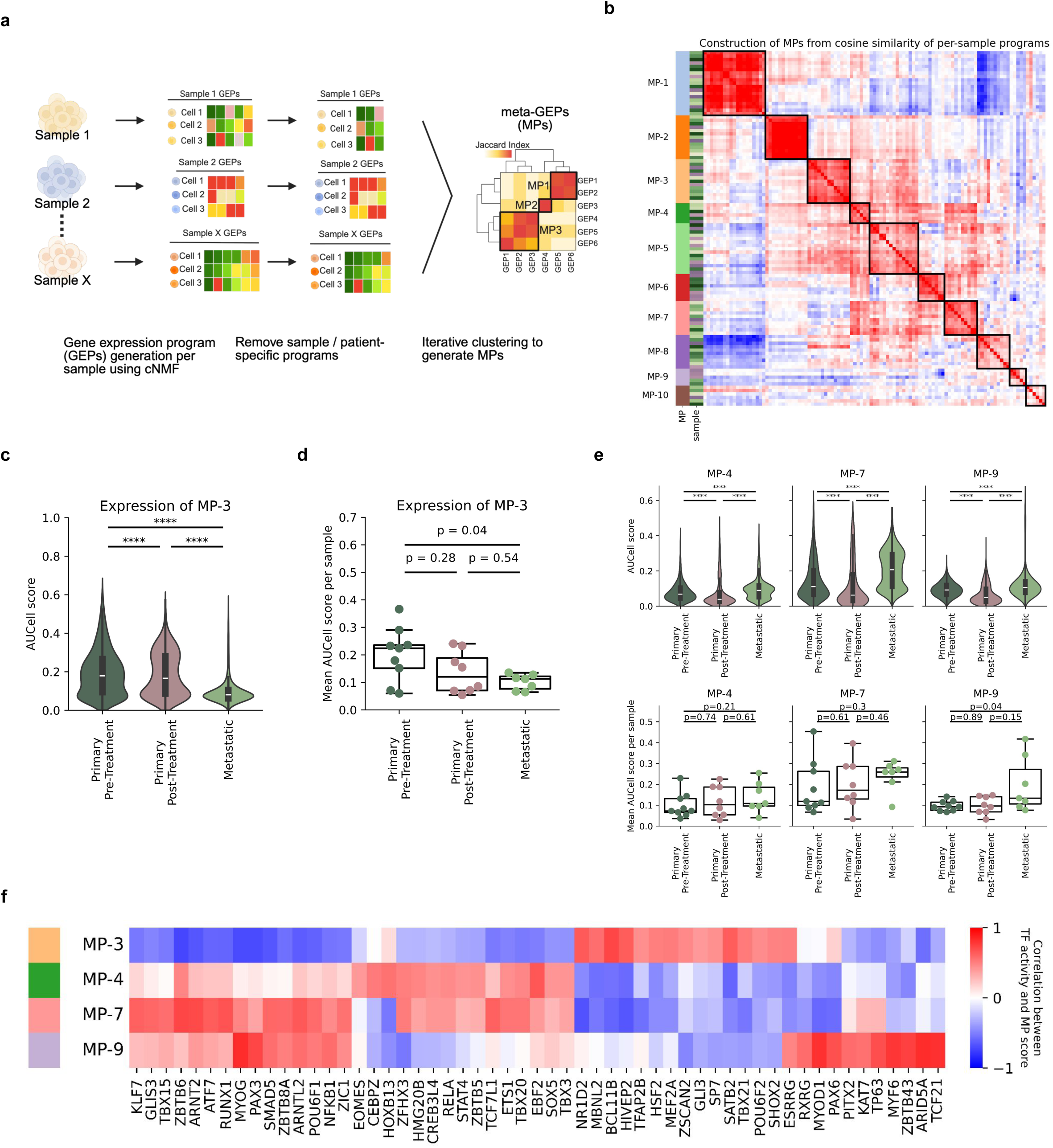
Localized and metastatic OS malignant cells adopt heterogeneous gene expression programs (a). Schematic detailing the computational workflow for deriving overarching meta-gene expression programs (MPs). Gene expression programs (GEPs) were initially inferred for each independent sample using consensus non-negative matrix factorization (cNMF). After filtering out patient-specific and redundant GEPs, the remaining GEPs were grouped via iterative clustering based on pairwise similarity (Jaccard index). **(b).** Heatmap of the Ward hierarchical clustering of individual sample-level GEPs into consensus MPs. The solid black boxes demarcate the 10 robustly defined MPs across the cohort. **(c).** MP-3 scored using AUCell on malignant cells stratified by primary-naive, primary-treated, and metastatic samples of origin. (MWU, **** denotes p < 0.0001), and **(d)** collapsed by mean per sample. **(e).** (Top) MP-4, MP-7, and MP-9 scored on AUCell on malignant cells stratified by primary-naive, primary-treated, and metastatic samples of origin. (MWU, **** denotes p < 0.0001), and (bottom) collapsed by mean per sample. **(f).** Heatmap of Pearson correlation coefficients between inferred transcription factor (regulon) activity scores and MP scores across all malignant cells. The matrix displays the top 15 most positively correlated regulons for each MP, highlighting putative transcriptional drivers of these specific cell states.

Based on the prior observation that osteoblastic marker gene expression was dampened in metastatic tumor cells **(Extended Data Fig. 1f)**, we used a comparative approach to more fully characterize differences in MPs between primary and metastatic tumor cells **(Extended Data Fig. 2b)**. The MP with the highest differential activity was MP-3, with significantly increased activity in primary naive and primary treated tumor cells relative to metastatic tumor cells **(Fig. 2c)**. In sample level analyses, we also observed a significant enrichment of MP-3 in primary naive samples relative to metastatic samples **(Fig. 2d)**. The MP-3 gene set included several genes classically associated with osteoblast differentiation, (e.g. *COL1A2, SP7, SATB2, and IBSP*) and its activity was positively correlated with expression of canonical osteoblastic genes **(Extended Data Fig. 2c-d)**. As such, these findings indicated that MP-3 likely represents an osteoblastic-like phenotype.

Among the remaining 9 MPs, we prioritized three additional MPs based on their differential activity between primary and metastatic tumor cells: MP-4, MP-7, and MP-9 **(Extended Data Fig. 2b)**. For each of these programs, we observed increased activity among metastatic tumor cells relative to primary naive and primary treated tumor cells **(Fig. 2e)**. While we were underpowered to observe differential activity for MP-4 and MP-7 at the sample level, we observed statistically significant enrichment of MP-9 among metastatic samples relative to the primary naive samples in our cohort **(Extended Data Fig. 2b)**, which was driven by high usage in 3 of 7 metastatic samples.

We inferred their putative biological functions by examining their constituent genes and correlating their expression in the OS malignant cells with existing pan-cancer metaprograms from the Curated Cancer Cell Atlas (3CA MPs)(24) **(Supplementary Table 4 / Extended Data Fig. 2e / Extended Data Fig. 2f)**. MP-4 included genes associated with less differentiated, non-lineage committed mesenchymal cell types such as *ENG, NRP1, COL4A1, and COL4A2*(26; 27; 28). MP-7 included non-lineage committed collagens (*COL3A1, COL6A1, COL6A2*(29; 30)), extracellular matrix genes (*POSTN* and *FN1*(31)), and the fibroblastic gene *ACTA2*(32). MP-4 and MP-7 were strongly positively correlated with the activity of epithelial-to-mesenchymal transition (EMT) 3CA MPs, further suggesting their association with a less, differentiated, non-lineage committed cell state. MP-9 included well established myogenic genes such as *RYR2*(33), *NEBL*(34), *MYH10*(35), and *FOXO3*(36) and AP-1 complex genes (*JUND, FOS,* and *FOSB*).

Given the strong activity correlation and gene composition overlap (27 genes shared) between MP-4 and MP-7, we sought to delineate their distinctive components **(Extended Data Fig. 2g)**. We found that the expression of genes unique to MP-4 correlated with Interferon/MHC-associated 3CA MPs, whereas MP-7 unique genes were broadly positively correlated with activity of EMT-related 3CA MPs **(Extended Data Fig. 2h)**. We observed that the expression of these MP-specific genes correlated strongly with each respective MP’s overall enrichment score **(Extended Data Fig. 2i)**.

We then inferred the activity of transcription factor (TF) regulons across the OS malignant cells **(Extended Data Fig. 2j)**, and identified distinct TF regulons whose activity correlated with each MP **(Fig. 2f)**. MP-3 was positively correlated with the activity of key osteoblast differentiation transcription factors, *SATB2*(37), *SP7* (38), and *SHOX2*(39). The activity of MP-4, MP-7, and MP-9 were negatively correlated with osteoblastic TF regulons, and instead were positively correlated with the activity scores of non-osteoblastic mesenchymal TFs such as those relevant to myogenic development (*PAX3*(40), *MYOG*(41)). MP-4 and MP-7 shared a significant proportion of positively correlated modules, including *RELA*, *TCF7L1*, and *TBX20*, TFs linked to maintaining a pre-lineage committed stem-like state, inflammation, cancer progression, and plasticity(42; 43; 44). The MP-9 program additionally showed strong positive correlation with the activity score of *MYF6* and *MYOD1*, well-established regulators of myogenesis, oncogenic plasticity, and limb development(45; 46).

In summary, we identified four OS MPs across OS tumor cells in our patient cohort with features of different mesenchymal lineages. While localized and metastatic tumor samples displayed activity of each of these MPs, MP-3 in particular harbored significantly higher activity in primary samples and was positively correlated with osteoblastic marker gene expression. Conversely, MP-4, MP-7, and MP-9 exhibited higher gradients of activity in metastatic tumor cells and were associated with non-osteoblastic mesenchymal gene expression.

### Modeling in primary human cells contextualizes OS MPs as mesenchymal programs with increased activity at different stages of osteoblastic differentiation

Given that the MPs observed in OS cells included genes and predicted TF regulons from a spectrum of mesenchymal cell states, we next sought to further contextualize these programs in the putative developmental context of OS, the MSC-to-osteoblast differentiation axis. We obtained primary human MSCs derived from discarded bone tissue of a healthy teenage patient undergoing orthopedic surgery and induced these MSCs to differentiate into bone-forming osteoblasts (OBs) by exposure to established OB differentiation conditions(47; 48) **(Fig. 3a)**. We characterized and benchmarked this experimental model of OB differentiation using a number of parallel approaches. We observed over a three week period that MSCs underwent expected morphological changes, deposited extracellular matrix, and mineralized that matrix consistent with an osteoblastic, bone-forming phenotype (Fig. 3b). Further, we found that both the canonical OB specifying TF, *RUNX2*, and the well established OB functional marker, *PHEX*, were upregulated over time in OB differentiation conditions **(Fig. 3c)**. We next characterized the expression changes induced in differentiating cells systematically by scRNA-seq. Due to the dense mineralized extracellular matrix deposited by mature osteoblastic cells, we could not robustly recover high quality single cells after 10 days of differentiation. We therefore used timepoints up to day 10 of differentiation in this system for scRNA-seq (Days 0, 3, 5, 7, 10). After processing, we obtained a dataset of 5,762 cells across all time points profiled from MSC to osteoblast differentiation **(Fig. 3d)**. We found that these cells broadly fell into three clusters: an MSC cluster and two differentiating clusters which we termed “early” and “intermediate” differentiating populations **(Fig. 3e)**. We annotated the clusters using known marker genes **(Fig. 3f)**. MSC markers (*ENG, THY1, NT5E*) remained stable, canonical osteoblast markers (*COL1A1, COL1A2*) progressively increased, and mature osteoblast markers (*ALPL, IBSP, BGLAP*) showed no significant upregulation. We confirmed this using a lineage agnostic trajectory inference tool, CytoTRACE2(49), which showed a progressive loss of potency from “multipotent” MSCs to predominantly “oligopotent” differentiating OBs, with a subset of more “unipotent” and “differentiated” cells in the intermediate differentiating cluster **(Fig. 3g)**. This suggested that our in vitro differentiation dataset spanned the spectrum of MSCs to immature osteoblastic/pre-osteoblastic cells, largely representing stem and early committed but non-terminally differentiated osteoblastic cell states.

**Fig. 3:**
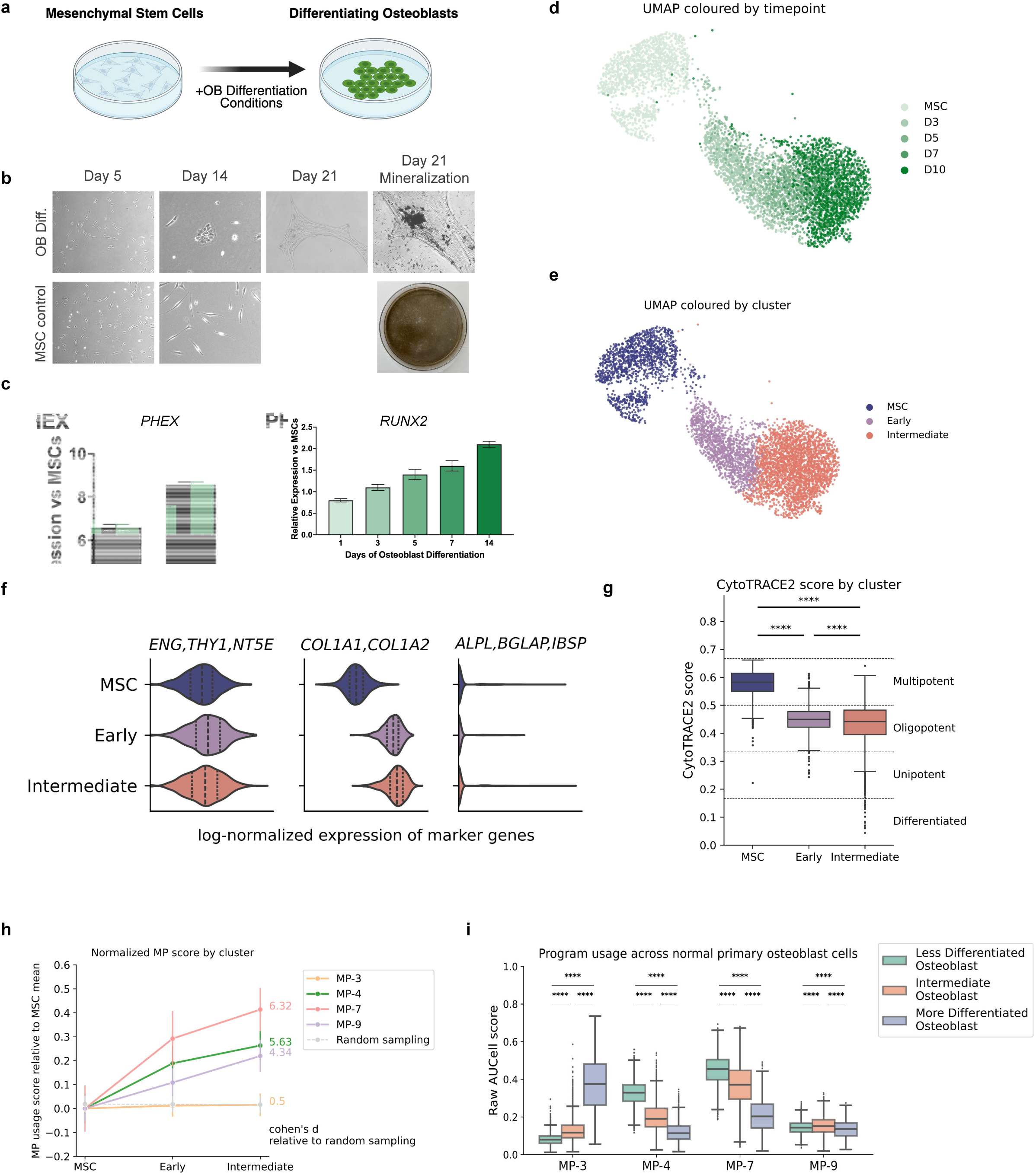
Tumor cell states reflect different stages of osteoblastic differentiation (a). Schematic detailing the differentiation protocol to model mesenchymal stem cells (MSC) to osteoblast. **(b).** Phase contrast images of differentiating cells at indicated time points. Right most image shows mineralization (Von Kossa) stains at Day 21 of differentiation. **(c).** qPCR for osteoblast markers *PHEX* and *RUNX2*. **(d).** UMAP visualization of the osteoblastic differentiation cells, colored by timepoint. **(e).** UMAP visualization of the osteoblastic differentiation cells, colored by differentiation stage clusters. **(f).** Normalized expression of marker genes across MSC (*ENG, THY1, NT5E*), osteoblastic lineage (*COL1A1, COL1A2*), and mature osteoblasts (*ALPL, IBSP, BGLAP*) by differentiation stage cluster. **(g).** cytoTRACE2 pseudotime distribution by differentiation stage cluster, with developmental potential categories as defined by cytoTRACE2. **(h).** Relative OS tumor MP activity across differentiation stages, normalized to their usage in MSC cells. As a control, we randomly sampled genes to use as a comparator of dynamic temporal expression across differentiation stages. We computed the cohen’s d for the deviation of the MP scores compared to the randomly sampled genes. **(i).** AUCell scores of MPs scored on non-malignant osteoblasts from tumor samples. We find that MP-3 is selectively expressed in more differentiated osteoblasts, while MP-4 and MP-7 are expressed significantly more highly in less differentiated osteoblasts (for all panels, **** denotes bonferroni corrected MWU p < 0.0001).

We next investigated whether the OS tumor-derived MPs were expressed in discrete phases of bone development **(Fig. 3h)**. We found that the osteoblastic MP-3 program, enriched in primary tumors, was lowly expressed in the stem and progenitor cells captured in our in vitro differentiation model. However, the other tumor programs MP-4, MP-7, and MP-9 increased in expression as MSCs transitioned to progenitor and non-mature osteoblastic cell types.

To evaluate the usage of these programs in more terminally differentiated cells, we assessed their expression in non-malignant osteoblasts identified in primary OS tumors which we stratified into less-differentiated, intermediate, and more-differentiated osteoblasts based on marker gene expression **(Extended Data Fig. 3a-b)**. We found that MP-3 was selectively enriched in the most differentiated osteoblasts, MP-4 and MP-7 were highly enriched in less differentiated osteoblasts and their expression decreased in more mature osteoblast populations, while MP-9 showed consistently low usage in all osteoblast subsets **(Fig. 3i)**. Taken together, these findings suggested that MP-3 was a mature osteoblastic program while MP-4, MP-7, and MP-9 were programs active in progenitor and immature osteoblastic cells that were suppressed during terminal differentiation.

### OS MPs are expressed in diverse mesenchymal cell types during embryonic development

Given that the OS MPs other than MP-3 were active in OB progenitor cells but deactivated during terminal OB differentiation, we next investigated the developmental context of OS MPs more broadly. First, we correlated MP usage in OS tumor cells with literature-derived markers of mature mesenchymal lineage cell types(18; 27; 45; 46; 50) **(Fig. 4a / Supplementary Table 3)**. As expected, MP-3 showed relatively strong positive correlation with OB marker gene expression and less strong positive correlation with chondrogenic marker genes. MP-4, MP-7, and MP-9 were positively correlated with mesenchymal stem cell (MSC) marker expression. MP-9 was also positively correlated with myoblast markers.

**Fig. 4:**
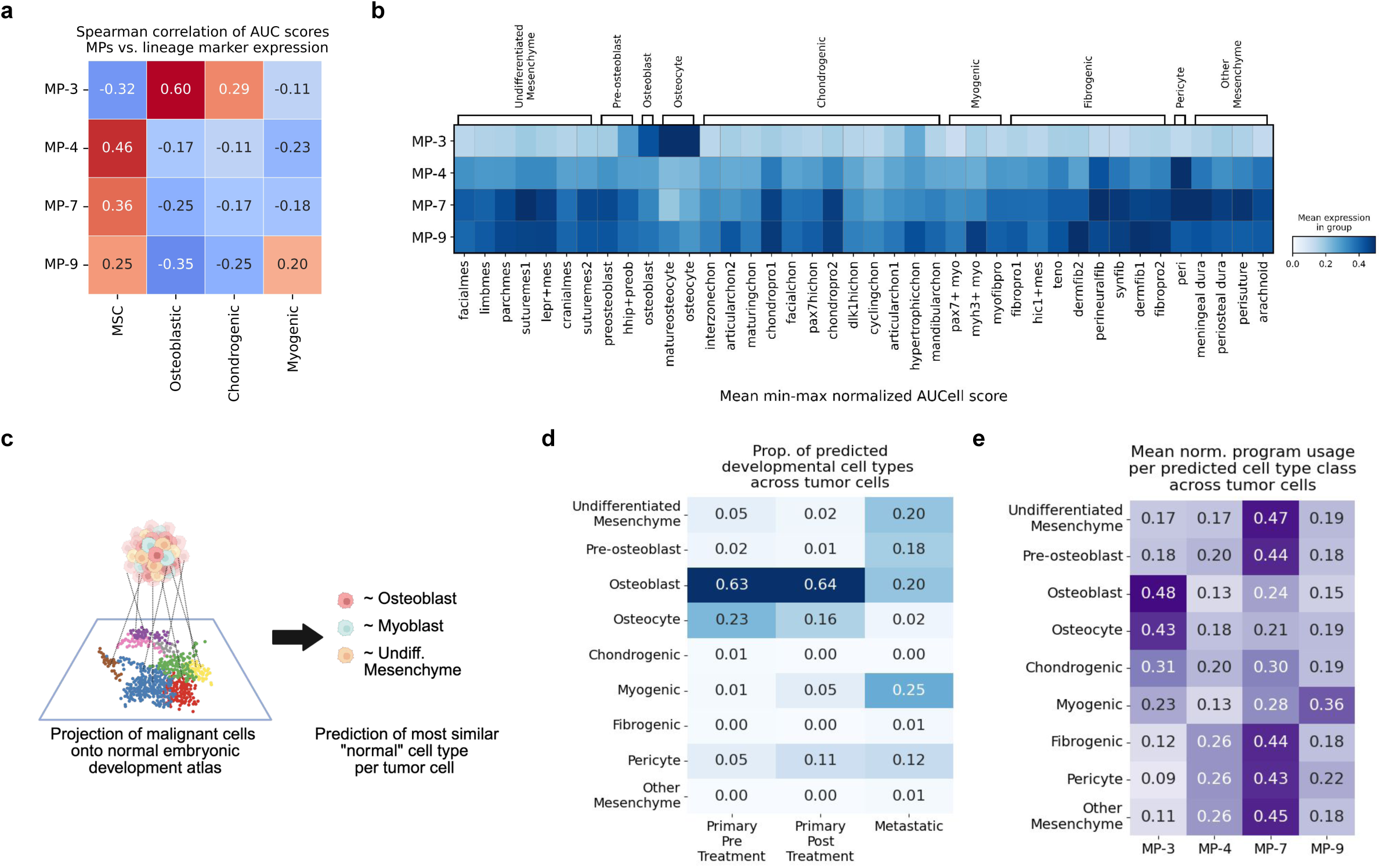
Tumor MPs are expressed in diverse mesenchymal cell types during embryonic development (a). Correlation of lineage-specific gene signature expression (chondroblast, osteoblast, MSC, myoblast) vs MP program scores across malignant cells. Each meta-program aligns with a distinct developmental lineage bias. **(b).** Mean min-max row normalized usage of the MPs per individual cell type annotated in the embryonic skeletal development atlas, grouped by broad cell class. **(c).** Schematic of training a semi-supervised model on the embryonic skeletal development dataset to predict the closest normal proxy of embryonic skeletal development dataset of each individual tumor cell. **(d).** Proportion of predicted cell type label projections using a semi-supervised model trained on the embryonic development dataset. We find that the primary (naive and treated) malignant cells are enriched for an osteoblastic-like phenotype, while metastatic malignant cells harbor broad heterogeneous, non-osteoblastic phenotypes (P < 0.0001, χ² test). **(e).** Distribution of the program usage by predicted normal proxy cell type. We first predict the normal proxy cell type for each individual tumor cell, and compute the mean score of each program for the cells predicted as that cell lineage. We find that the MP-3 program is used in tumor cells that are predicted to be osteoblastic-like, while MP-4 and MP-7 programs are used in undifferentiated or other mesenchyme-like tumor cells, MP-9 are used additionally for myogenic-like tumor cells.

To further define MP developmental contexts, we evaluated their expression across all mesenchymal cell types in a single cell atlas of embryonic skeletal development(51) **(Extended Data Fig. 3c / Fig. 4b)**. Consistent with our prior observations, MP-3 usage was highest in mature osteoblasts and osteocytes and relatively flat in other cell types. MP-4, MP-7, and MP-9 all showed usage in a broad range of mesenchymal cell types including undifferentiated mesenchyme. MP-4 and MP-7 showed high usage in pericytes. MP-7 and MP-9 showed high usage in chondrocytes and fibroblastic cell types. MP-9 was unique in its high usage in myoblasts. These findings indicated that the OS primary tumor enriched MP-3 was an expression program relatively restricted to mature osteoblastic cell populations while MP-4, MP-7, and MP-9 were expressed in a diverse spectrum of normal mesenchymal cell types with skewing toward particular cell populations.

To characterize OS cell states independent of MP usage, we utilized a single-cell label transfer method to project malignant OS cells directly onto this atlas **(Fig. 4c)**. Consistent with the enrichment of MP-3, we found that the majority of primary tumor cells (86% treatment naive, 80% treated) mapped onto normal osteoblasts or osteocytes. In contrast, metastatic OS cells mapped to a diverse spectrum of primitive, non-osteoblastic cell states, including myogenic (25%), undifferentiated mesenchyme (20%), and pre-osteoblast (18%) populations **(Fig. 4d)**, consistent with a recurrent pattern of phenotypic divergence and lost osteoblastic lineage fidelity during metastatic progression. To further assess the connection between cell state and OS MPs, we analyzed MP usage in OS cells stratified by cell type labels transferred from the developmental atlas **(Fig. 4e)**. We found that OS cells resembling normal osteoblasts and osteocytes highly expressed MP-3 with relatively low usage of the other programs. MP-4, MP-7, and MP-9 were expressed across a broad range of cell types. MP-7 expression skewed toward undifferentiated mesenchyme-like and pre-osteoblast-like cells while MP-9 expression skewed toward myogenic-like cells. Collectively, these findings suggested that primary OS tumor cells predominantly adopt an osteoblast-like cell state, while metastatic OS tumor cells adopt non-osteoblastic cell states similar to a broad spectrum of other mesenchymal cell types.

### Transcriptional state in metastatic OS evolves independently from genetic clonality

To directly evaluate transcriptomic evolution between primary and metastatic tumors, we compared the usage of the tumor MPs across two primary-metastatic tumor pairs **(Fig. 5a)**. Consistent with our whole cohort analysis, we found that metastatic tumors exhibited decreased usage of the MP-3 program compared to their primary tumor of origin. Increased expression of non-osteoblastic MPs also occurred in each metastatic tumor, although the predominant MP differed between patients. In the patient 15 pair, MP-7 was the most highly expressed in metastatic tumor cells, while MP-9 predominated in metastatic cells from patient 17. When we projected cells from these pairs onto normal mesenchymal cell types, we found that the majority of primary tumor cells in both samples (88% in patient 15, 64% in patient 17) projected onto osteoblasts, while metastatic cells projected onto mesenchymal cell types distinct from their paired primary tumors **(Extended Data Fig. 4a)**. In the patient 15 pair, 42% of metastatic cells projected onto undifferentiated mesenchyme compared to 5% of primary tumor cells. In the patient 17 pair, 83% of metastatic cells projected onto myogenic cell types, compared to 0% of primary tumor cells.

**Fig. 5:**
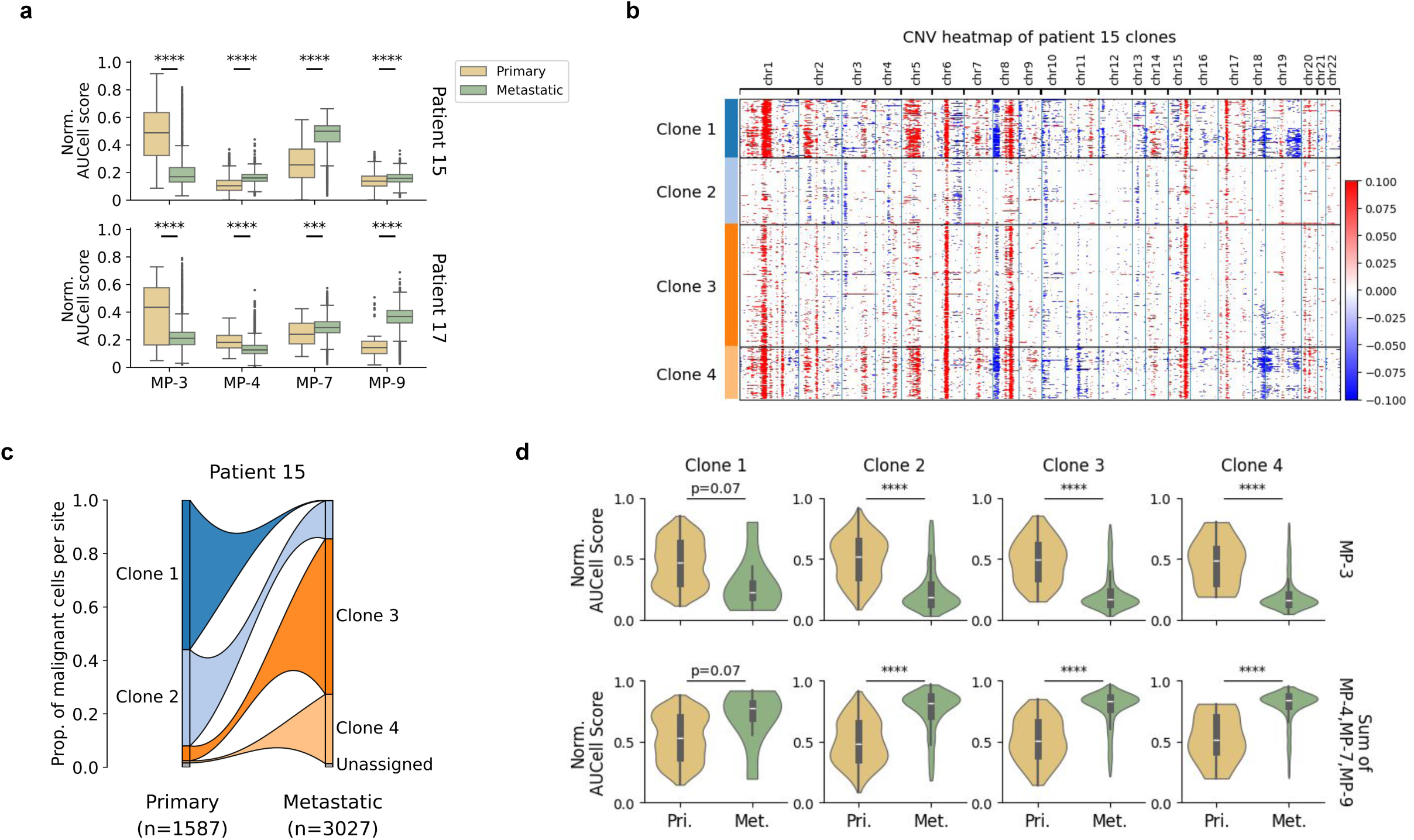
Metastatic subclones arise in primary tumors and adopt site specific transcriptional states independent of genetic clonality (a). Distribution of program scores between paired samples per patient (Patients 15 and 17). We observe depletion of MP-3 at the metastatic site, and an increased usage of a metastatic-enriched program (MP-7 and MP-9) in each of the pairs. (Bonferroni corrected MWU, **** denotes p < 0.0001) **(b).** Inferred copy number profiles for malignant cells in the paired samples for patient 15, clustered by predicted subclone. **(c).** Proportion of malignant cells assigned to each clone in the primary and metastatic samples for patient 15. **(d).** Distribution of program usage within each subclone in primary and metastatic sites for patient 15.

To investigate how genetic clonal evolution related to this observed transcriptional evolution, we next inferred distinct genetic subclones in primary-metastatic tumor pairs by inferring copy number alterations based on scRNA-seq profiles **(Extended Data Fig. 4b-c / Fig. 5b / Methods)**. In patient 15, we assigned 98.5% of tumor cells to four distinct clones **(Fig. 5c)**. For patient 17, we identified clonal cell populations but had insufficient clone representation in the primary tumor sample to robustly associate cell state with clonality across all clones, likely due to a limited number of malignant cells. Comparing clone representation between primary and metastatic sites in patient 15, we observed that all four clones were present in the primary and metastatic tumors, indicating that at this resolution all identified genetic clones had the capacity to metastasize. Further, we observed that two identified clones (Clones 1/2) contracted while the other two clones (Clones 3/4) expanded in the metastatic tumor **(Fig. 5c)**. It is unclear if these changes in the relative representation of each clone were due to differential metastatic fitness or sampling bias.

We next sought to evaluate the variability in MP usage across sites within subclonal populations of tumor cells. By comparing MP scores between the primary and metastatic tumors for each clone, we observed that primary tumor clones exhibited high usage of the osteoblast-like MP-3 and low usage of non-osteoblastic MPs (MP-4, MP-7, and MP-9) **(Fig. 5d)**. This pattern reversed across all clones at the metastatic site, regardless of whether the clone had expanded or contracted, suggesting the transcriptional state of malignant cells evolved during metastatic progression, independent of genetic clonality. Furthermore, we observed that intra-clonal transcriptional diversity was highly constrained at the metastatic site **(Fig. 5d)**. Quantifying this phenotypic narrowing using Euclidean distance revealed a tighter distribution of per-cell MP usage around the clonal mean in metastatic samples compared to primary tumors **(Extended Data Fig. 4d)**. Collectively, these findings suggest the metastasis bottleneck may impose a selective pressure on OS cells, driving them to decrease osteoblast-like expression and uniformly enrich other mesenchymal MPs.

### Enrichment of non-osteoblastic mesenchymal transcriptional states in localized post-treatment OS tumors

There is a growing body of evidence linking therapy resistance to metastatic progression across tumor types(52; 53; 54; 55). To control for inter-sample heterogeneity, we subsequently focused on the two available paired samples (patients 12 and 18) to better understand the characteristics of post-treatment disease. Similar to the metastatic setting, we observed a relative loss of osteoblastic MP usage (MP-3) and an increase in other mesenchymal MP (MP-4, MP-7) usage in the post-treatment samples **(Fig. 6a)**. Filtering our projections onto the embryonic skeletal development atlas for the malignant cells in these paired samples, we found that pre-treatment cells largely projected onto osteoblasts and osteocytes while post-treatment cells projected onto a broader spectrum of mesenchymal cell types **(Fig. 6b)**. These findings indicate that the OS malignant cells that persist at the primary tumor site after multi-agent chemotherapy are enriched for non-osteoblastic mesenchymal cell states, paralleling our observations in metastatic disease.

**Fig. 6:**
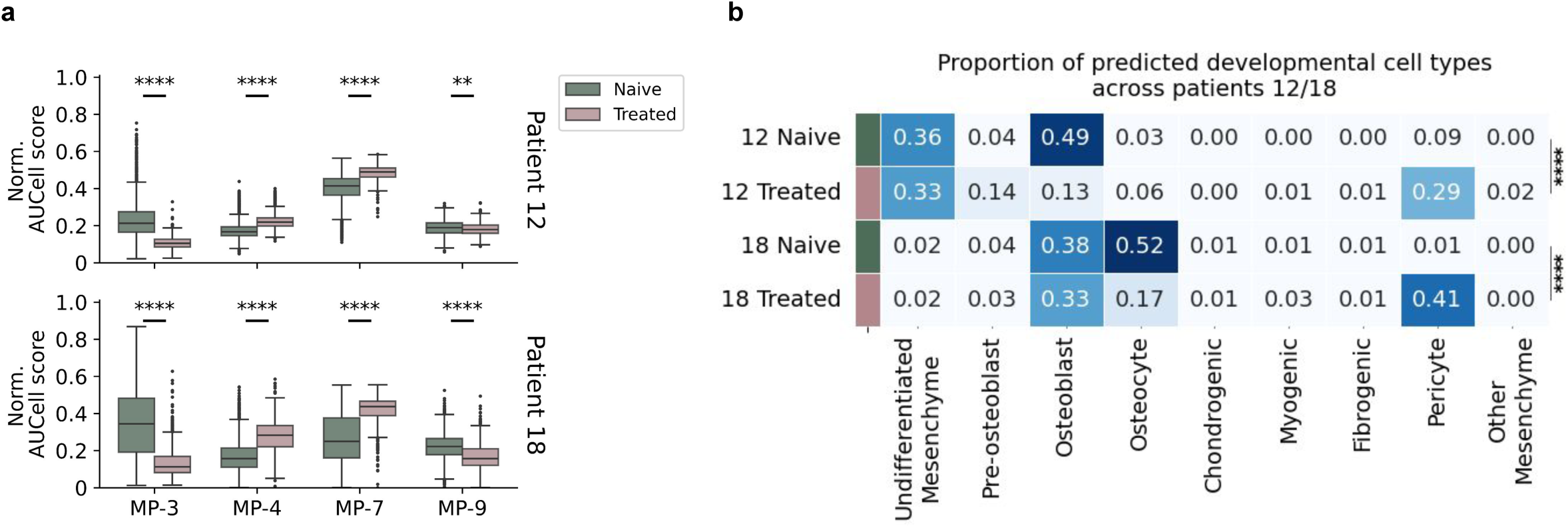
Non-osteoblastic tumor programs are enriched in post-treatment localized tumors (a). Distribution of cell-normalized program scores between paired pre-post treatment localized samples per patient (Patients 12 and 18). We observe depletion of the canonical osteoblast-like program Pri-1 (MP-3) in the post-treatment sample, and increased usage of the MP-4 and MP-7 programs in each of the pairs. (Bonferroni corrected MWU, ** denotes p < 0.01, **** denotes p < 0.0001) **(b).** Proportion of predicted proxy cell type projections from the embryonic skeletal development dataset-trained model. We find that the pre-treatment malignant cells are enriched for an osteoblastic-like phenotype, while post-treatment malignant cells harbor broad heterogenous, non-osteoblastic phenotypes (P < 0.0001, *χ*^2^ test per-patient).

## Discussion

In this study, we discovered diverse mesenchymal gene expression programs across localized and metastatic OS patient tumors. Most notably, we observed the loss of an osteoblastic transcriptional phenotype in metastatic tumor cells, a feature that may reflect a key adaptive property of metastatic fitness in OS.

We defined tumor metaprograms (MPs), and identified a primary tumor-enriched MP (MP-3) along with three other highly active mesenchymal MPs (MP-4, MP-7, MP-9). We found that utilization of MP-4, MP-7, and MP-9 was heterogeneous within and between metastatic tumor cell populations, underscoring the heterogeneity of OS and indicating potentially distinct pathways to metastatic success. Through regulon analysis we identified TFs that may drive the acquisition of non-osteoblastic cell phenotypes and confer metastatic fitness. By integrating tumor data with data from an *in vitro* system modeling early osteoblastic differentiation, and a published atlas of embryonic skeletal development, we found that MP-3 represented a program uniquely active in mature osteoblasts and osteocytes. By contrast, expression of the MP-4, MP-7, and MP-9 programs increased during early osteoblastic differentiation, but was lost in mature osteoblasts, suggesting that these programs are active in progenitor cell types. Our findings were thematically concordant with recent studies that have identified distinct early and late osteoblast-like cell states in OS and associations with distinct clinical features and outcomes(21; 56; 57). Further, these MPs were significantly enriched in other mesenchymal cell populations including undifferentiated mesenchyme, myogenic cells, and pericytes indicating that OS cells, particularly in metastasis, activate non-osteoblastic programs.

By evaluating paired primary and metastatic samples, we observed a transcriptional shift at the sample and subclonal levels: primary tumor cells predominantly expressed the osteoblastic program, but activated non-osteoblastic, mesenchymal transcriptional programs at the metastatic site. These transcriptional shifts occurred intra-clonally, irrespective of whether the clone expanded or contracted, suggesting that cell state change away from the osteoblastic lineage is an adaptive metastatic trait in OS. This supports a model of transcriptional plasticity in OS metastatic progression, rather than selection of rare, transcriptionally static subpopulations of malignant cells. Additionally, this adaptive plasticity appears to link the selective pressure of metastatic colonization with that of treatment effect. Comparing naive and treated primary tumor pairs, we observed a marked loss of the osteoblastic program (MP-3) usage and a relative expansion of malignant cells with mesenchymal program usage (MP-4, MP-7) among OS cells persisting after multi-agent chemotherapy. When projected onto normal cells, these treated OS cells were found to have lost their osteoblastic identity and occupy a broad spectrum of mesenchymal states. These data suggest that therapeutic pressure may select for subpopulations of surviving cells in the highly plastic states required for metastasis, highlighting that the transcriptional adaptations required to survive therapeutic stress overlap with those that confer fitness at the metastatic site.

While this study provides new insight into the transcriptional underpinnings of metastatic progression in OS, certain limitations must be acknowledged. Given the epidemiological rarity of OS and its substantial inter-tumoral heterogeneity, we were underpowered to capture the full breadth of relevant transcriptional states driving OS metastasis, for which larger patient cohorts are necessary. In addition, our analysis on pre-/post-treatment and primary-metastatic pairs was limited, and larger cohorts are required to pinpoint drivers of treatment resistance and disease progression. Our clonal inference was based on transcriptomically-inferred copy number alteration profiles, which do not have the resolution of DNA-based clonal tracing. Finally, further mechanistic work is necessary to delineate the tumor cell intrinsic and extrinsic mediators of the transcriptional cell states identified in this study.

Our study represents a significant step forward in understanding the differential biology of localized and metastatic OS. Our findings support a model in which metastatic OS cells have higher activity of non-osteoblastic mesenchymal gene programs, which may endow them with the necessary properties for invasion, colonization, and survival. While the maintenance of a canonical osteoblastic transcriptional state in primary tumor cells appears to be less permissive to dissemination and colonization of distant tissues, it may allow persistence in the tissue of origin. These findings resonate beyond OS, suggesting a shared paradigm of metastatic progression in aggressive pediatric sarcomas and other solid tumors, with recent studies suggesting context-specific aberrant developmental drivers in other tumor types(58; 59). Together, these insights provide a foundation to inform novel therapeutic strategies that look beyond optimization of bulk tumor eradication to specifically target the gene expression programs and regulatory networks that enable and maintain highly plastic, less-differentiated cell states that fuel metastatic progression.

## Methods

### Osteosarcoma patient tumor samples

We obtained 28 frozen tissue specimens from pediatric patients (age range 4-21) with a diagnosis of osteosarcoma (OS) that were seen at Dana-Farber/Boston Children’s Cancer and Blood Disorders Center collected with written informed consent and ethics approval by the Dana-Farber Cancer Institute Institutional Review Board under protocol number 17-104. This cohort included primary, localized tumors (18 samples), and metastatic tumors (10 samples). All available clinical characteristics of these patient samples are summarized in (Supplementary Table 1).

Of note, one case was omitted from analysis due to a revision of pathologic diagnosis to “pleomorphic high-grade sarcoma” on confirmatory pathology review, and after quality control of the sequencing data, we removed one additional poor quality sample, defined by having <100 nuclei retained. In addition, we did not confidently identify any malignant cells in samples 19P and 19M based on their expression-inferred copy number alteration profiles. We retained these normal samples in the overall dataset cell type annotation and tumor microenvironment analyses. As such, 24 samples were used for downstream malignant cell analyses in this study.

### Sample preparation, cell isolation, and library preparation for tumor single-nuclei transcriptomic sequencing

Nuclei isolation was performed as previously described (60), with minor modifications. In short, low-retention microcentrifuge tubes (Fisher Scientific, Hampton, NH, USA) were used throughout the procedure to minimize nuclei loss. Frozen tissue was mechanically dissociated by pipetting and homogenized in TST solution plus 0.3% Tween-20 and RNAase inhibitors, filtered through a 30 um MACS SmartStrainer (Miltenyi Biotec, Germany), and pelleted by centrifugation for four minutes at 500 x g at 4C. The nuclei pellet was resuspended in 200ul of 1x PBS, 1% BSA, and RNAase inhibitor, and nuclei were counted by eye using INCYTO C-Chip Neubauer Improved Disposable Hemacytometers (VWR International Ltd., Radnor, PA, USA).

Approximately 8,000 nuclei per sample were loaded per channel of the Chromium Next GEM Chip K for processing on the 10x Chromium Controller (10x Genomics, Pleasanton, CA, USA) followed by cDNA generation and library construction, as per manufacturer’s instructions (Chromium Next GEM Single Cell 5’ Reagent Kits v2 User Guide, Rev E). Libraries were normalized and pooled for sequencing on NovaSeq SP-100 flow cells (Illumina, Inc., San Diego, CA, USA) using run parameters 26, 10, 10, 90.

### Single-nuclei RNA sequencing data processing

The generated sequencing data was obtained in a raw base call (BCL) file format, which was then processed using the cumulus implementation of the 10X Genomics CellRanger toolkit (version 6.0.1) to generate gene expression counts (61; 62). In this pipeline, “CellRanger mkfastq” with default parameters was first run to generate FASTQ files from BCL files. Then, the “CellRanger count” step was used to align the reads to the 10X Genomics GRCh38-2020-A (GENCODE v32 / Ensembl v98) reference genome, including the intronic reads, with an expected cell count of 5,000, to generate per-cell barcode-unique molecular identifier (UMI) count matrices for each sample (10x Genomics, Technical Note – Interpreting Intronic and Antisense Reads in Single Cell Gene Expression Data, Rev A).

We next filtered each individual sample’s count matrix for potential ambient RNA using CellBender (version 0.3.0) (63). We first ran the computational pipeline on default settings, and manually reviewed the results to remove likely empty droplets. After filtering out potential ambient RNA, we filtered out low quality cells using scanpy (version 1.9.8) (64) built-in functions. Here, we defined “low quality” nuclei as those with less than 300 genes or 500 UMI counts expressed, or more than 6000 genes and/or 50,000 UMI counts expressed. Additionally, we filtered out nuclei that had no ribosomal gene expression and more than 20% and 5% of mitochondrial and hemoglobin genes over total expressed genes respectively. We also removed any sparsely expressed genes (genes expressed in less than 5 cells in each sample), all mitochondrial and hemoglobin genes, and *MALAT-1*, a non-coding gene known to be highly detected independent of protocol in 10X Genomics sequencing from further downstream analysis (65). Next, we removed potential doublets from the filtered count matrix using scrublet (version 0.2.3) (66).

### Normalization, clustering, and visualization of snRNA-seq data

Various downstream analyses and data visualization required normalization and clustering of the snRNA-seq counts matrices. Log-normalization was performed by normalizing counts to the median total counts or to 10,000 total counts, followed by log(x+1) transformation. To identify biologically highly variable genes independent of technical variance, we adapted a Pythonic implementation of the scran modelGeneVar function from the voyager package (version 0.1.1) (67; 68). We used 3000 highly variable genes as default. Clustering was performed by first Principal Component Analysis (PCA) on the highly variable genes identified, constructed a nearest neighbor graph, and then Leiden clustering(69). Visualizations of snRNA-seq data were generated using Uniform Manifold Approximation and Projection (UMAP) with an effective minimum distance between embedded points of 0.5, using the computed nearest neighbor graph and partition-based graph abstraction (PAGA) as input(70).

### Cell identity gene expression programs to annotate cell types

To annotate nuclei for their cell identity across the dataset at a per-nuclei resolution, we performed consensus non-negative matrix factorization (cNMF) (version 1.3.0)(71) for each per-sample site (eg. bone, lung) to generate “cell type identity” gene expression programs (GEPs), which we used to annotate the nuclei(72). We performed this on each sample site to ensure samples were annotated consistently across tumor samples, given that tumor microenvironment cells from similar biological sites are known to have a similar gene expression profile(73).

For each run, the 3000 highly variable genes (HVGs) were first computed from the coding genes. In addition to these HVGs, we manually included a subset of literature-derived cell type marker genes to ensure inclusion of markers for less common cell types (10). cNMF was run on these gene lists for 200 iterations across a broad range of programs generated (“K” = 2-50). We manually selected the optimal K independently for each run, as determined by the silhouette score. Next, we annotated each gene expression program based on a list of canonical cell type marker genes (Supplementary Table 3). We then aggregate the usage of programs that were annotated as the same cell type to compute a “cell type identity usage” score for each cell type, and normalize the score to sum to 1 for each cell. We annotated each cell with the cell type it had highest usage for, but annotated cells with no dominant program present (cell type identity usage score *<* 0 5) as “low confidence”. The “low confidence” cells were annotated by performing leiden clustering, identifying the cell types it clustered with.

We confirmed the robustness of these annotations by confirming consistency of cell type annotations between sample sites, and performed multiple rounds of manual revision and adjustment of any annotation discrepancies. We removed any cells that had inconsistent annotations, or expressed mixed lineages. We benchmarked this against other approaches to evaluate the suitability of this approach, and evaluated its sensitivity (Supplementary Material 1).

### Malignant cell annotation

After we annotated all cells for their corresponding “normal” cell type classes, we used inferCNVpy (version 0.4.5) to annotate the malignant cells in each sample (inferCNV of the Trinity CTAT Project). Given the variable proportion of immune cells in each sample, we first generated a uniform “normal cell” reference dataset by randomly sampling 20% from all immune (Myeloid, NK/T, Osteoclast, B) cells available across the cohort (3,327 cells). We used this common reference as the “control” for all per-sample inferCNVpy runs, and the “case” for each run was defined as all mesenchymal lineage cells in each sample. While the lineage of origin for OS is thought to be osteoblastic, all mesenchymal cells were included as “cases” to capture malignant cells with more heterogeneous expression profiles. inferCNVpy was run on the log-normalized gene expression matrix using a sliding gene window of 250 genes, a CNV score log-fold change clip of 1, and a step of 1 (computing all possible windows). Then, leiden clustering on the cells’ CNV profiles was conducted on a neighborhood graph computed with 20 neighbors and 20 PCs at a clustering resolution of 1. Otherwise, default parameters were used. This approach clusters cells based on their inferred copy number alteration profile, and computes a “CNV score” per cluster.

We defined malignant cells as cells that were assigned a CNV score higher than one standard deviation above the mean CNV score of the control immune cell population. This approach was taken as OS does not have a ubiquitously recurring copy number alteration event. We manually reviewed the inferCNV plot for each sample to confirm the presence of broad copy number alterations in the cells annotated as malignant, and performed manual adjustments on the CNV cluster annotations accordingly.

### Gene signature scoring

We used the decoupler (version 1.6.0)(74) implementation of AUCell(75) and the scanpy(64) score_genes function to score various gene signatures, such as gene expression programs and pathway gene sets.

### Statistical Enrichment

In this study, we used different categorical (eg. Leiden clustering) and continuous (eg. program scoring) variables to characterize and stratify scRNA-seq data. In order to evaluate whether these variables are enriched / biased towards any class (eg. sample, cell type), we utilized a number of statistical metrics. Statistical tests are described where used, using Mann-Whitney U (MWU) tests as default for continuous variables.

### Lineage agnostic/aware evaluation of cell differentiation

CytoTRACE2, a lineage agnostic measure of differentiation based on transcriptional diversity, was used to predict degree of cell differentiation(49). In addition, we scored malignant cells using a mesenchymal lineage aware metric of differentiation, expanding an approach used to study rhabdomyosarcoma, a pediatric sarcoma arising from the myogenic lineage(58). In brief, we scored each cell for mesenchymal stem cell (MSC), osteogenic, myogenic, and chondroblastic lineages using manually curated lists of canonical marker genes for each lineage (Supplementary Table 3). We normalized these scores to estimate each cell’s differentiation status across the mesenchymal landscape.

### Malignant cell state metaprogram generation

To better characterize the cellular states broadly observed within and across all samples, we performed consensus non-negative matrix factorization (cNMF) (version 1.3.0)(71) on each individual sample’s tumor cells, and then conducted iterative supervised clustering of all per-sample generated programs to generate meta-gene expression programs (MPs)(23; 25).

cNMF was run for 200 iterations on 3000 highly variable genes, and to minimize biased selection of variables, we selected a broad range of values (3,4,5,6,7,8,9,10) for the number of components generated (“K”). Of note, we confirmed and included additional known mesenchymal lineage marker genes, for a total of 3018 highly variable genes, adapting an approach previously described(8). For each sample, we selected the optimal K based on manual selection of the K with the highest stability.

To cluster and generate MPs, we removed all low prevalence programs. Here, we defined low prevalence as being the predominantly expressed program (normalized usage of *≥* 0 5) in less than 10 cells within the sample. Next, we initially Ward clustered all retained programs using their gene weights to summarize and generate recurrently expressed “MPs”. The clustering was then manually reviewed and individual programs with low correlation were further removed. For each MP, we defined the representative gene list by reranking the constituent programs’ mean gene weights, and taking the top 100 genes as its gene set. Through this iterative clustering approach, we generated robust, consensus “MPs” that broadly describe recurrently represented cell states present across the cohort.

### Regulon activity inference

pySCENIC (version 0.12.0) was used to infer the activity of known transcription factors in the malignant cells(75). We ran the program per sample using default parameters, using the default motif (2022 SCENIC+ human motif collection) and search space input (10 kilobases around the transcription start site (TSS), and 500 base pairs upstream / 100 base pairs downstream of the TSS) files provided with the pySCENIC package, and a list of known transcription factors(76; 77). After the per-sample runs, we identified robustly expressed regulons by filtering for regulons found to be “active” in multiple samples and are found to have moderate expression (mean AUCell Score > 0.001 across the cohort). The per-cell activity of the regulons were evaluated using the AUCell scores computed by the pipeline.

### Mesenchymal Stem Cells

Cells were initially isolated from a discarded normal bone fragment from a pediatric patient undergoing orthopedic surgery with approval from the Institutional Review Board of the Centre Hospitalier Universitaire Vaudois (CHUV, University of Lausanne). The sample was de-identified and transferred to DFCI in accordance with Institutional Review Board approval under a non-human subjects designation.

### MSC-to-Osteoblast Differentiation

Prior to differentiation, MSCs were maintained in Iscove’s modified Dulbecco’s medium, 10% FCS, and 10 ng/mL platelet-derived growth factor BB (PeProtech EC) as previously described(47). Osteoblastic differentiation was induced as previously described(48). Cells were plated at a density of 3×103 cells/cm^2^ in MSC media. The following day media was removed and replaced with osteogenic media: DMEM, 10% FBS, 100nM dexamethasone, 0.05mM L-ascorbic acid 2-phosphate, 10mM *β*-glycerophosphate which was replaced every 3-4 days.

### Osteoblast Marker Gene Expression Analysis

Cells were lysed in 1 ml TRIzol reagent (Life Technologies), and RNA was extracted with 200 *µ*l chloroform. The organic phase was isolated, and 700 *µ*l of ethanol was added. RNA was purified from this solution using the RNeasy Micro Kit (Qiagen). cDNA was synthesized using the High Capacity RNA-to-cDNA kit (ABI) according to the manufacturer’s protocol. Marker gene expression was assessed by RT–qPCR analysis for ALPL and RUNX2 using optimized TaqMan Gene Expression Assay primers and TaqMan Gene Expression Master Mix (Life Technologies). The probes used for the RT–qPCR are as follows: ALPL, Hs01029144_m1; RUNX2, Hs01047973_m1; GAPDH, Hs02758991_g1.

### Mineralization (Von Kossa) Staining

Cells were washed with PBS then fixed for 30 minutes using 10% formalin. Following fixation cells were washed x3 with deionized water then incubated with 2% sodium nitrate in the dark for 10 minutes. Plates were thoroughly washed with deionized water and exposed to UV light for 20 minutes.

### MSC-to-Osteoblast differentiation scRNA-sequencing

Undifferentiated MSCs and MSCs grown in osteoblast differentiation conditions for 3, 5, 7, or 10 days were processed on the same day in parallel to mitigate batch effects. 5000 cells from each condition were then processed for scRNA-seq using the PIP-seq T2 3’ Single Cell RNA Kit v4.0PLUS based on a particle-tempted emulsification method(78) following the manufacturer’s instructions. Following library preparation and manufacturer suggested QC, samples were sequenced on an Illumina NextSeq 500 instrument on the same flowcell.

### *in vitro* MSC-to-Osteoblast scRNA-seq data processing

Single-cell RNA sequencing of the in-vitro MSC-to-osteoblast differentiation system sequencing was performed using PIP-seq v4 chemistry (Fluent BioSciences). Raw sequencing reads were processed using PIPseeker (v3.0.5) running the full pipeline. Reads were aligned to the GRCh38 reference genome (PIPseeker reference release 2022.04) using the bundled STARsolo aligner with the GeneFull feature, which counts both exotic and intronic reads.

Individual sample count matrices were merged then processed as described in “Single-nuclei RNA sequencing data processing” and “Normalization, clustering, and visualization of snRNA-seq data” section, with minor changes. Initially, cells were filtered to retain only those expressing at least 2,000 unique genes. To exclude low-quality or damaged cells, we filtered cells with more than 10% of mitochondrial and 20% ribosomal gene expression over total expressed genes. For initial quality control, we calculated the log2(TPM+1) for all genes and retained genes with an average expression of *≥* 4. Scrublet and top 0.5% of cells with gene and UMI counts were used to filter potential doublets. PCA was performed using 40 PCs and 10 neighbors, and Leiden clustering was performed at 0.3 resolution. After clustering, we removed poor quality clusters, which we defined as clusters with median gene count <3000 and median mitochondrial gene content >7%, and a cell cycle cluster, identified by scoring for cell cycle phases. Gene program activity was scored using all of the genes available in the data matrices, irrespective of the filtering performed.

### Embryonic skeletal development snRNA-seq atlas

We obtained the processed count matrices from a recent embryonic skeletal development atlas(51). Briefly, these single-nuclei RNA-seq atlases were generated from healthy human embryonic tissue collected from elective termination procedures at different timepoints to model skeletal cell-fate determination through development. We used the published cell type annotations provided with the count matrices as described in the original publication, and filtered for mesenchymal lineage cells.

### Normal developmental atlas label transfer

We adapted a previously described approach(79) to project tumor cells onto their transcriptomically closest normal development system cell type labels. In brief, we used the scANVI method in scArches, a semi-supervised machine learning approach that is initially trained on the annotated normal “reference” atlas, and then we use the trained model to predict malignant cell’s closest developmental phenotype(80). The models were trained for 100 epochs using default parameters. Cells were assigned to the cell type they had the highest predicted score for.

### Clonal inference

To estimate clonal evolution across pairs of samples from the same patient at different clinical timepoints (eg. Primary - metastatic), we extend the same approach as described in the “Malignant Cell Annotation” section, clustering on the predicted copy number alteration profiles across all of the malignant cells in the paired samples. Cells that were assigned to the same cluster were annotated as the same clone, and these profiles were manually confirmed using the visualized copy number alteration profiles.

### Visualization and formatting

We used CoMut to visually present the clinical characteristics of the patient tumor sample cohort(81). All graphical schematics used in figures were generated using BioRender [Biorender.com]. The color palettes were adapted from the GitHub MetBrewer repository [github.com/BlakeRMills/MetBrewer]. All other visualization tools used are described in the manuscript’s GitHub repository. LLMs, such as Google Gemini, were used to improve language and readability of the text throughout, which was then manually reviewed and edited further.

## Article Information

### Resource Availability

#### Lead Contact

Further information and requests for data, resources and materials described in this manuscript should be directed to, and will be fulfilled upon reasonable request by co-corresponding author, Riaz Gillani, M.D.

#### Materials Availability

This study did not generate any new materials or reagents.

#### Data Availability

The raw sequencing data for the generated patient tumor snRNA-seq and in-vitro normal osteoblast differentiation scRNA-seq will be made available at [dbGaP: phs004090.v1.p1](https://dbgap.ncbi.nlm.nih.gov/) upon final publication. Supplementary data and code used in this study’s computational analysis is publicly available on GitHub at https://github.com/gillanilab/metastatic-osteosarcoma-singlecell.

A processed version of the patient tumor snRNA-seq data generated as part of this study is available through the Alex’s Lemonade Stand Foundation, single cell Pediatric Cancer Atlas (ALSF scPCA), accession number SCPCP000017 (https://scpca.alexslemonade.org/projects/SCPCP000017). The data available on this portal has been processed using the ALSF scPCA computational pipeline, independent of the processing described in this manuscript, and is detailed in the data portal’s documentation(82). In this manuscript, we provide a comparison between the data processing performed in this study and the ALSF scPCA data in Supplementary Material 4 to highlight differences.

## Acknowledgments

We gratefully acknowledge the patients, their families and caregivers, and hospital personnel, without whom this study would not have been possible. We also thank the members of the Gillani, Bernstein, and Van Allen Labs at the Dana-Farber Cancer Institute for their thoughtful comments and feedback throughout this study. The generation of patient tumor snRNA-seq data was funded by a ALSF single-cell Pediatric Cancer Atlas grant (N.B.C.). This work was supported by Alex’s Lemonade Stand Foundation (ALSF) Young Investigator Awards (YIA) (J.J.M, R.G.); Damon Runyon-St. Jude Pediatric Cancer Research Fellowship (J.J.M.); American Society of Clinical Oncology (ASCO) Career Development Award (R.G.); ASCO-Conquer Cancer Sarcoma Foundation of America YIA (R.G.); Boston Children’s Hospital (BCH) Translational Research Program Mentored Translational Investigator Service Award (R.G.); Dana-Farber Cancer Institute (DFCI) Wong Family Award in Translational Oncology (R.G.); DFCI-BCH Pedals for Pediatrics grant (R.G., K.A.J.); Rally Foundation Career Development Award (R.G.); Sarcoma Foundation of America Jay Vernon Jackson Memorial Research Award (R.G.); Department of Defense Peer-Reviewed Cancer Research Program Fellow Career Development Award CA220721 (R.G.); Break Through Cancer’s Defying Osteosarcoma Project (J.J.M, R.G., K.A.J.); Count Me In’s Osteosarcoma Project (R.G., K.A.J.); DFCI-BCH Pan-Mass Challenge, Midnight Riders and Precision for Kids Teams (K.A.J.); ALSF Pediatric Oncology Student Training Program (Grant Award-1480465, funded by Northwestern Mutual) (L.H.G.); Innovation in Cancer Informatics award (E.M.V.A.); Ellison Foundation (E.M.V.A.); Ambrose Monell Foundation (E.M.V.A.); DFCI Richard and Nancy Lubin Family Endowed Chair (B.E.B.); American Cancer Society Research Professor (B.E.B.); St. Jude Research Collaborative (B.E.B); Harvard Ludwig Center (B.E.B.).

This work was additionally supported by NIH grants U2CCA252974 (R.G.); K08CA276701 (R.G.); Pediatric Oncology T32 (5T32CA136432-15) (J.J.M.); R50CA265182 (J.P.), R01CA227388 (E.M.V.A.); R37CA222574 (E.M.V.A.); U01CA233100 (E.M.V.A.); U2CCA233195 (E.M.V.A.). The content is solely the responsibility of the authors and does not necessarily represent the official views of the National Institutes of Health (NIH).

## Contributions

Conceptualization: N.B.C., R.G., J.J.M., B.E.B., Y.T.; Methodology: Y.T., J.J.M., R.G.; Investigation: all authors; Formal analysis and visualization: Y.T., J.J.M., R.G.; Funding acquisition: J.J.M., R.G., N.B.C., B.E.B.; Supervision: R.G., B.E.B.; Manuscript - original draft: Y.T., J.J.M., R.G., B.E.B.; Manuscript - review and editing: all authors.

---

**Extended Data Fig. 1:**
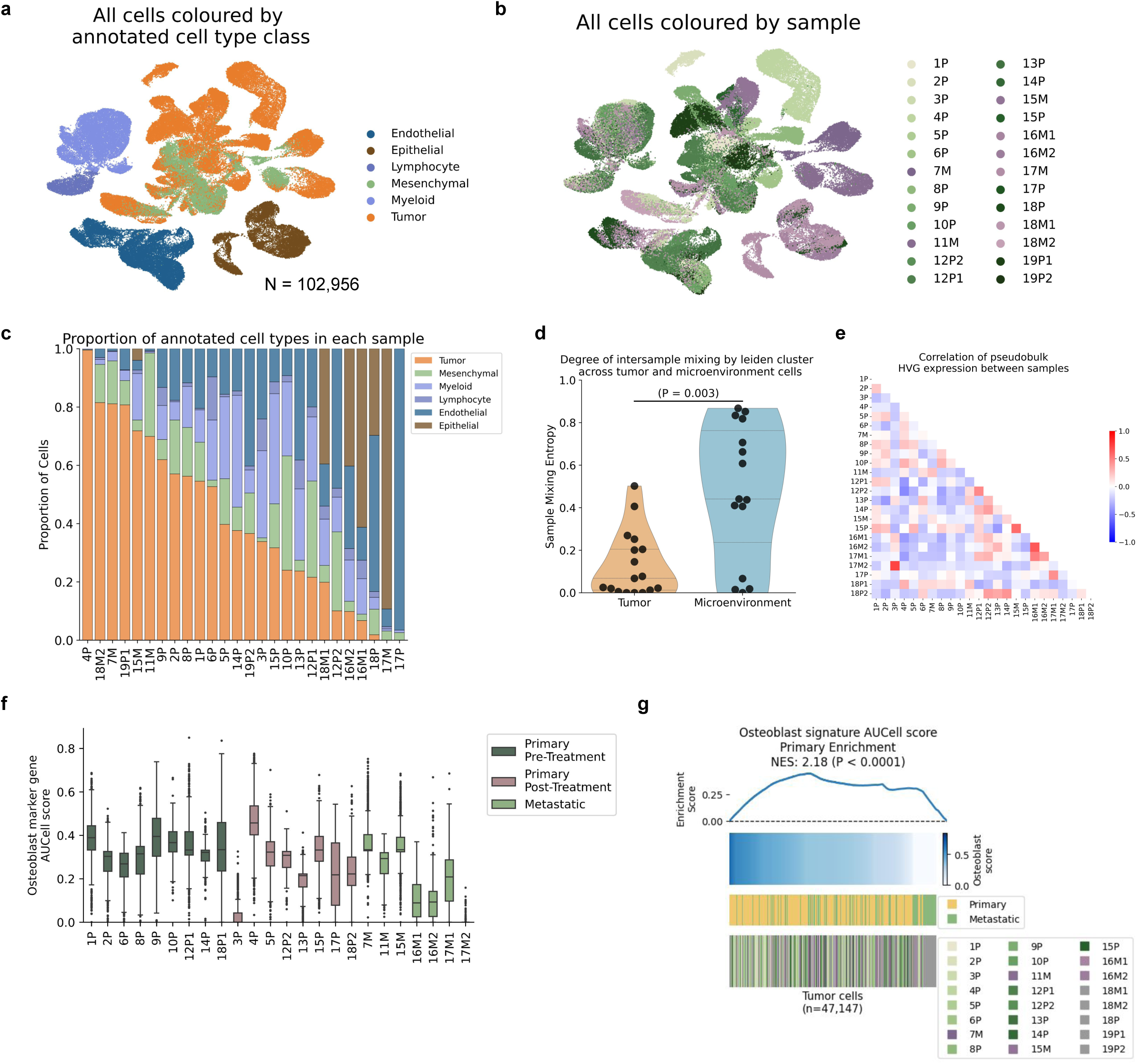
Overview of snRNA-seq dataset (a). UMAP projection of the complete snRNA-seq dataset (n = 102,956 nuclei), colored by broad cell type class (endothelial, epithelial, lymphocyte, myeloid, mesenchymal, tumor). **(b).** UMAP of all cells, colored by sample of origin, showing dataset composition across 26 patient samples. **(c).** Proportion of annotated cell types within each tumor sample. **(d).** Violin plots comparing sample-mixing entropy between tumor and tumor microenvironment clusters, revealing significantly different inter-sample mixing (P < 0.001, Mann-Whitney U Test), indicative of patient-specific transcriptional expression in tumor cells. **(e).** Heatmap showing correlation between expression of the cohort’s top 3,000 highly variable genes across malignant cells from individual samples. Low inter-sample correlation suggests high inter-tumoral heterogeneity. **(f).** Distribution of Osteoblast signature AUCell score by sample, categorized by primary-naive, primary-treated, and metastatic tumor of origin. **(g).** Ranked distribution of Osteoblast signature AUCell score (bottom) between primary and metastatic tumor of origin. NES (normalized enrichment score) was calculated for enrichment in primary tumors.

**Extended Data Fig. 2:**
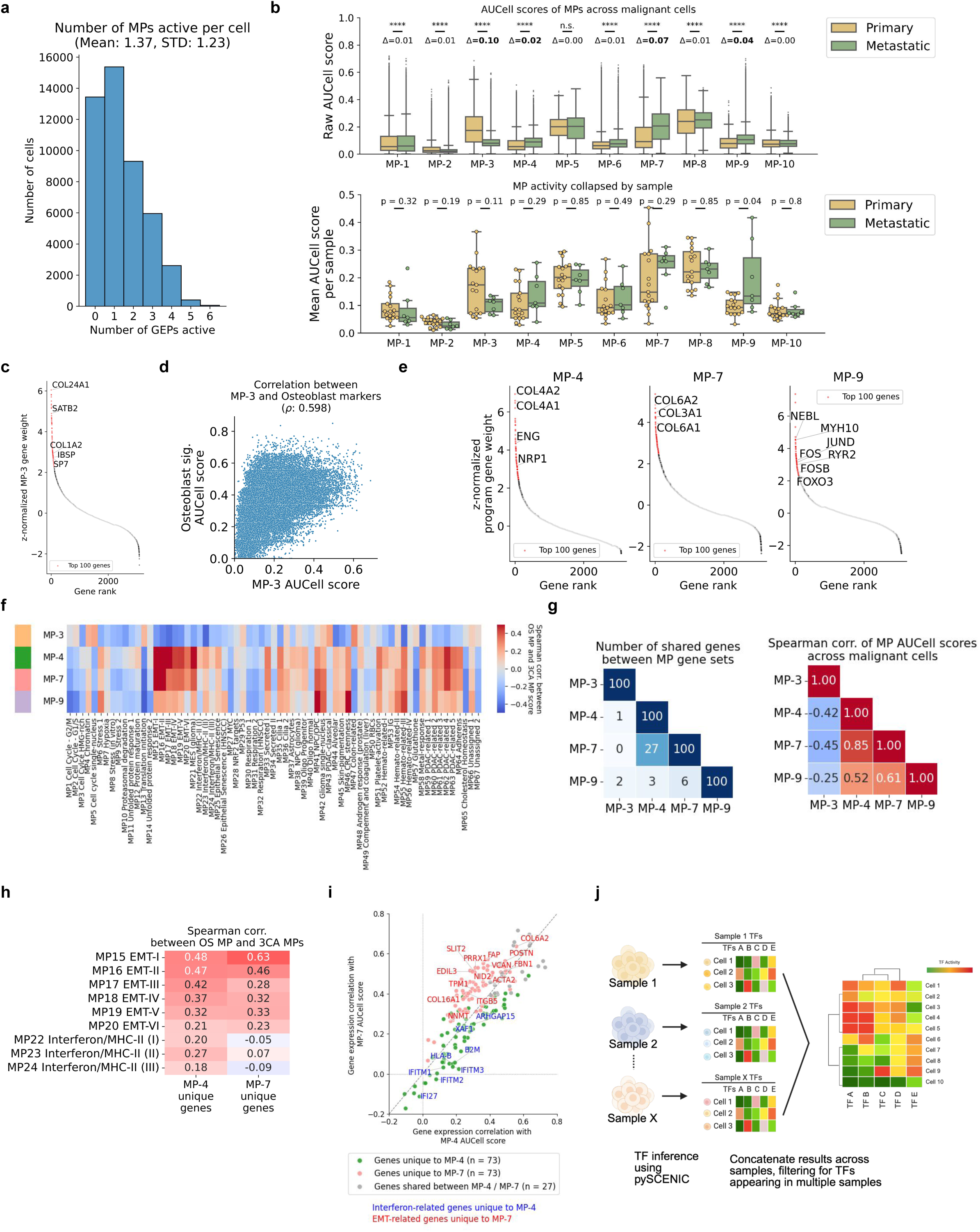
Generation and annotation of MPs derived from malignant cells (a). Distribution of the number of active MP programs per cell. We defined active as a z-score normalized MP score > 1, finding that a mean of 1.37±1.23 programs are active across malignant cells. **(b).** MP scores for malignant cells (top) and collapsed mean values per sample (bottom) stratified by primary versus metastatic sample origin. Statistical tests were performed using Bonferroni-corrected MWU tests (* denotes p < 0.05, ** denotes p < 0.01). **(c).** Rank of genes by z-normalized contribution weights to MP-3. Top 100 genes are highlighted in red. **(d).** Spearman correlation between the AUCell scores of MP-3 and the Osteoblast marker gene signature scores. **(e).** Rank of genes by z-normalized contribution weights to MP-4, MP-7, and MP-9. Top 100 genes are highlighted in red. **(f).** Spearman correlation between the MP scores and the Curated Cancer Cell Atlas (3CA)27 MPs scored across the malignant cells. **(g).** (Left) Number of shared genes between MPs. We find that there is a significant number of genes (27) that overlap between MP-4 and MP-7. (Right) Spearman correlation between the activity of MPs. We also find that the activity of MP-4 and MP-7 are correlated. **(h).** Spearman correlation between the EMT/Interferon-related 3CA MPs and the activity of the genes unique to MP-4 and MP-7 (n = 73 genes each) scored across the malignant cells. **(i).** Correlation between the MP-4 and MP-7 and the expression of the MP genes across the malignant cells. We highlighted the Interferon/EMT-related genes unique to MP-4 and MP-7 respectively, and found that while the genes were positively correlated with both program usage, but skewed towards each respective program. **(j).** Graphical schematic of transcription factor (TF) activity inference from scRNA-seq data. We run pySCENIC on all malignant cells per sample, and aggregate results, filtering for robustly expressed TFs found across multiple samples.

**Extended Data Fig. 3:**
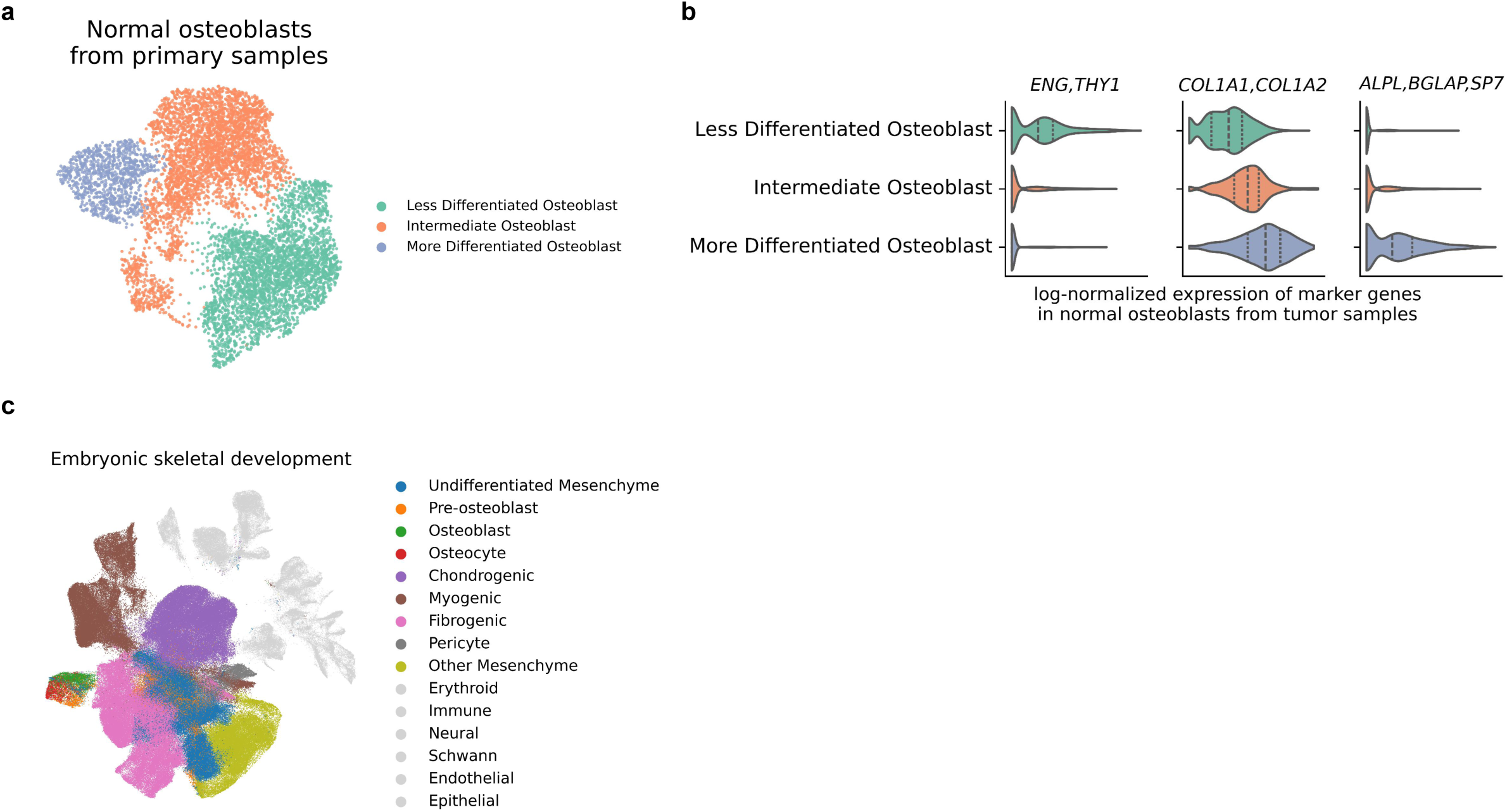
Contextualization of the normal osteoblastic and mesenchymal differentiation datasets (a). UMAP visualization of the non-malignant osteoblast cells from the primary tumor samples, annotated by differentiation status. **(b).** Stratification of the non-malignant osteoblasts in the primary tumor samples was performed using canonical marker genes, across the three subtypes identified - less differentiated osteoblasts (MSC marker genes: *THY1, ENG*), intermediate osteoblasts (Osteoblast marker genes: *COL1A1, COL1A2*), and more differentiated osteoblasts (Mature osteoblast marker genes: *ALPL, IBSP, SP7*). **(c).** UMAP visualization of the overall embryonic skeletal development, colored by cell type lineage, highlighting the mesenchymal cell type lineages.

**Extended Data Fig. 4:**
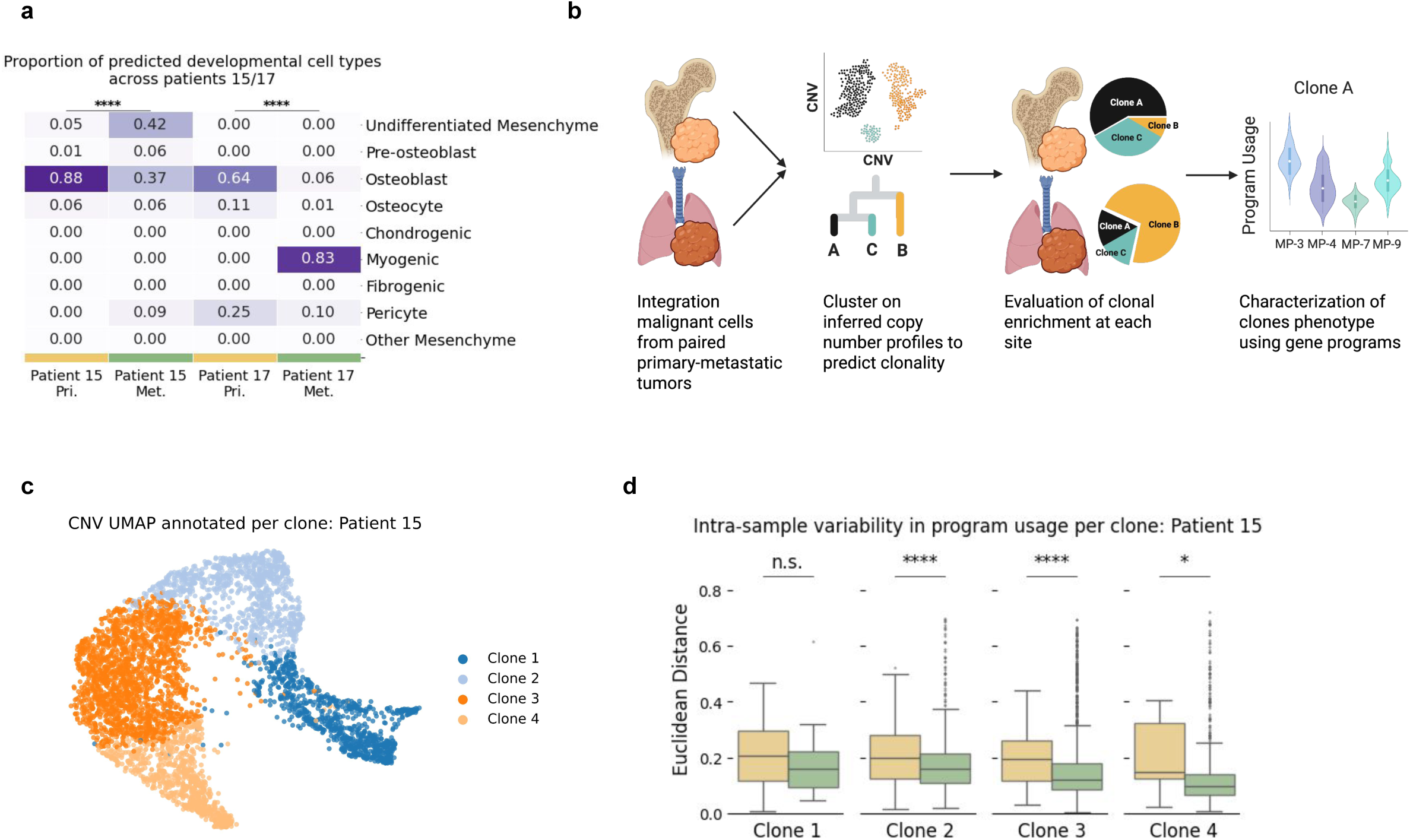
Clustering of inferred copy number alteration profiles enables clonality prediction (a). Proportion of predicted proxy cell type projections from the embryonic skeletal development dataset-trained model. We find that the primary malignant cells are enriched for an osteoblastic-like phenotype, while metastatic malignant cells harbor broad heterogenous, non-osteoblastic phenotypes (P < 0.0001, χ² test per-patient). **(b).** Schematic of method to predict clonality from inferred copy number alteration profiles. **(c).** UMAP visualization of clustering performed on the inferred copy number alteration profiles from patient 15 paired samples (sample 15P/15M), colored by predicted clones. **(d).** Distribution of euclidean distance, a statistical proxy for transcriptional heterogeneity, between program usage per cell in subclones stratified by primary and metastatic sites. We find that within each clone, the heterogeneity of program usage decreases at the metastatic site.

