## Supplementary Materials for "Metastatic Osteosarcoma is Characterized by Loss of Osteoblastic Lineage Fidelity"

#### Supplementary Material 1. cNMF-based cell type annotation

In single-cell studies, cell types are commonly annotated by utilizing community detection methods such as Leiden and Louvain clustering to generate cell clusters, and identifying the highly expressed or differentially expressed marker genes in each cluster(Traag et al.; Wolf et al.). However, as cell type annotation is performed on a per-cluster resolution in these approaches, it is prone to noise and potential misannotation of rare/transitory cell types that may cluster with more abundantly found cell types. In this study, we used cNMF(Kotliar et al.), a matrix decomposition method, to infer the gene expression programs (GEPs) dominantly present across the dataset, annotate individual cells for cell types using the GEPs robustly utilized in each cell, and refined these annotations iteratively **(Methods)**.

We evaluated the sensitivity of this method to several manually selected parameters, using the largest run (bone samples) as a representative example. In this study, we elected to use k=18 GEPs from the cNMF output. We first evaluated the variability in generated GEPs by selecting other "k" values with high stability (k = 16-20) **(Supp. Fig. 1a)**. We found that the generated programs were highly consistent; 14 programs were conserved (>0.75 Jaccard index of top 100 genes) across all 5 selected "k"s, and an additional 4 programs were conserved across at least 4 "k"s **(Supp. Fig. 1b)**. Next, we annotated each generated GEP using marker genes and calculated a "cell type score" for each cell by summing the weights of programs annotated to the same cell type. Each cell was assigned the cell type for which it scored highest. Cells with a cell type score < 0.5 for all categories were annotated as "low confidence" (5,105 cells; 7.14% of total) **(Supp. Fig. 1c)**. Analysis of these "low confidence" cells revealed that a significant proportion (78.9%) were annotated as "mesenchyme" or "malignant" after all annotation steps **(Supp. Fig 1d)**. Collectively, these results demonstrate that our approach yields consistent annotations despite variability in parameter selection.

We further compared this approach against standard methods, such as annotating individual clusters per sample site using canonical marker genes. We found a high rate of agreement (93.0%) between the GEP-based and clustering-based methods **(Supp. Fig 1e)**. Most discrepancies were driven by cells that the GEP-based method filtered out due to mixed expression of marker genes **(Supp. Fig 1f)**. While both methods broadly agreed, the GEP-based method was more accurate in filtering out doublet-like cells.

#### Supplementary Material 2. Granular tumor microenvironment annotation

This study focused on intrinsic OS tumor cell plasticity. Recognizing the nature of the newly generated snRNA-seq dataset, we performed cell type annotation on each broad cell type class (Endothelial, Epithelial, Immune, and Mesenchyme) to generate a broad overview of the tumor microenvironment in this cohort.

***Endothelial.*** We identified five major endothelial subtypes across primary and metastatic OS samples: arterial, capillary, venous, lymphatic, and tip-like endothelial cells described in prior studies **(Supp. Fig 2a)**. Proportional analysis revealed shifts in endothelial composition between primary and metastatic samples, consistent with vascular remodeling in metastatic niches.

***Epithelial.*** Epithelial cells were classified into alveolar (type 1 and 2), ciliated, proximal and distal tubular (including thick ascending limb), and intercalated cells **(Supp. Fig 2b)**. Notably, smaller populations of cells in bone-localized samples were annotated as epithelial-like and expressed canonical alveolar markers (*AGER*, *PDPN*); these were provisionally annotated as "alveolar" despite their atypical location. As expected, metastatic samples exhibited a higher abundance of epithelial cells, consistent with the lung/kidney tissues of origin.

***Mesenchymal.*** Mesenchymal populations were categorized into cancer-associated fibroblasts (CAFs), mesenchymal stem cells (MSCs), osteoblasts, pericytes, and myoblasts **(Supp. Fig 2c)**. Notably, primary tumors were enriched for osteoblast-lineage cells, as expected of bone tissue, while metastatic sites harbored an increased abundance of myoblast-like populations. CAFs were consistently present in both compartments. These shifts may reflect both developmental reprogramming and stromal adaptation to metastatic microenvironments.

***Immune.*** The immune compartment consisted of diverse lymphoids **(Supp. Fig 2d)** and myeloid populations **(Supp. Fig 2e)**. Lymphoid populations comprised CD4+, CD8+, regulatory, and exhausted T cells, as well as NK cells, B cells, and plasma cells. A small population of lower-quality T cells lacked sufficient resolution for subclassification. Myeloid cells included tumor-associated macrophages (TAMs), monocytes, myeloid-derived suppressor cells (MDSCs), dendritic cells (cDC1, cDC2, pDCs), mast cells, and osteoclasts. Immune composition varied between primary and metastatic tumors, with relative increases in exhausted T cells and monocyte/MDSC populations, and decreases in macrophage and CD4+ T cell populations at the metastatic site.

#### Supplementary Material 3. Description of all generated MPs

In this study, we generated meta-gene expression programs (MPs) from patient tumor derived single-nuclei RNA-seq data. The approach taken is described in the Methods, and we focused on a subset of these MPs (MP-3, MP-4, MP-7, MP-9) in the primary study. Here, we have provided a broad description of the remaining 6 MPs (MP-1, MP-2, MP-5, MP-6, MP-8, MP-10) identified across the tumor data that we do not further characterize in the primary study. The full list of top constituent genes for each MP is provided in **(Supplementary Table 4)**. We extended much of the computational analysis performed in the primary study to characterize the programs and reviewed their constituent genes to annotate and group the programs.

***MP-1 and MP-10.*** These MPs are correlated with each other (Supp. Fig. 3a), and are distinctly anti-correlated with the broader mesenchymal and osteoblastic MPs. They are also both uniquely correlated with stress and respiration related 3CA MPs (Supp. Fig. 3b). In line with their constituent genes, we define MP-1 as a Stress-related MP (including *HSP90AB1*, *EEF1A1*, *EEF2*), and MP-10 as a tRNA / metabolism related MP (including *IARS*, *GARS*, *HSPA9*).

***MP-2.*** We annotated as a cell cycle program, given its inclusion of canonical G2/M phase and mitotic spindle markers (*ASPM*, *CENPE*, *BUB1B*, *TOP2A*), its strong correlation to the G2/M phase cell cycle signature score **(Supp. Fig. 3d)**, and its direct alignment with the Cell Cycle G2/M 3CA MP **(Supp. Fig. 3b)**.

***MP-6.*** In the primary analysis, we described the non-osteoblastic, mesenchymal-like phenotypes of MP-4 and MP-7 in detail. We observed substantial overlap and correlation between MP-4, MP-6, and MP-7, both in terms of their constituent genes and their activity AUCell scores **(Supp. Fig. 3a)**. We characterized MP-6 as an additional mesenchymal MP that includes robust mesenchymal markers (*PDGFRB*) and structural collagens (*COL18A1*, *COL6A2*, *COL3A1*), and is only positively correlated with the MSC signature score **(Supp. Fig. 3e)**.

***MP-5 and MP-8.*** Our primary MP analysis identified distinct non-osteoblastic MPs, such as MP-9 described. In addition to MP-9, we find that MP-5 is characterized by neuronal development genes (*TENM3*, *NAV3*, *SEMA3A*) and is positively correlated with the usage of TF regulons such as *FOXP1*, *MEF2C*, and *PBX3* **(Supp. Fig. 3c)**. We additionally identified MP-8 that included similar cell adhesion and synaptic organization genes (*DLG2*, *FAT3*, *RORA*). Interestingly, we find that these MPs are correlated with MP-3 **(Supp. Fig 3a)**, an MP we annotated as canonically osteoblastic, but its usage across the embryonic skeletal development atlas **(Supp. Fig 3f)** suggests that these transcriptional programs are non-specific to osteoblastic lineage cell types.

#### Supplementary Material 4. Comparison between alevin-fry and 10x CellRanger

A processed version of the snRNA-seq data is available through the Alex's Lemonade Stand Foundation's single cell Pediatric Cancer Atlas (ALSF scPCA - accession number SCPCP000017). Raw FASTQ files generated by the 10x Genomics "mkfastq" step were submitted to ALSF for inclusion in their repository. The ALSF scPCA uses a data processing pipeline independent of the one used in this study(Hawkins et al.), utilizing alevin-fry (a pseudoalignment-to-transcriptome approach) rather than the 10x Genomics CellRanger pipeline used here. We performed a quantitative comparison at each step of data processing available through ALSF scPCA, comparing the ALSF scPCA's unfiltered, filtered, and processed matrices to our pipeline's post-CellRanger, post-CellBender, and post-QC matrices, respectively **(Supp. Fig 4a)**. We used the 27 samples that passed QC in both pipelines for this comparison.

***Unfiltered vs post-CellRanger.*** We first evaluated the differences in the genes in both reference genomes used. The alevin-fry reference (Ensembl 104) was substantially larger than the CellRanger reference (GRCh38-2020-A/Ensembl 98), containing 60,319 unique gene IDs compared to 36,601, with an intersection of 36,456 IDs (Jaccard Index = 0.603). Notably, the alevin-fry reference included 20,003 IDs without gene names, and 26.7% ± 16.1% of total reads in the alevin-fry data were assigned to these unnamed IDs **(Supp. Fig 4b)**. Additionally, 5.84% ± 3.44% of alevin-fry reads were assigned to genes entirely absent from the CellRanger reference, a marked contrast to the CellRanger data where only 0.049% ± 0.018% of reads mapped to genes missing from the alevin-fry index **(Supp. Fig 4c)**. Comparison of the matrices revealed a high correlation in the number of expressed genes (0.899). However, the number of nonempty cells showed a very low correlation (0.118), and the normalized Jaccard Index for exact cell overlap was poor (median < 0.3) **(Supp. Fig 4d)**. In total, we found significant discrepancy between the reference genome, and what each pipeline defined as an unfiltered / raw-read.

***Filtered vs post-CellBender.*** There was a marked improvement in concordance at this stage **(Supp. Fig 4e)**. The correlation for the number of genes rose to 0.946, and the correlation for the number of cells increased significantly to 0.898. The normalized Jaccard Index for cell overlap also improved, indicating much stronger agreement between the pipelines after removing empty droplets.

***Processed vs post-QC.*** We observed the highest concordance at this final step **(Supp. Fig 4f)**. The correlation for the number of genes reached 0.961, while the number of cells maintained a strong correlation of 0.823. The normalized Jaccard Index for exact cell overlap remained high, at a similar distribution to that of the prior step.

While the overlap between retained genes and cells increased with the amount of processing, we found significant differences between these two common approaches. The most significant divergence stemmed from the reference genomes used, though differences persisted at each step. These analyses highlight the inter-pipeline processing variations that arise despite identical sequencing data; these should be considered and adjusted for during experimental design, particularly when integrating multiple cohorts.

#### Supplementary Figure 1

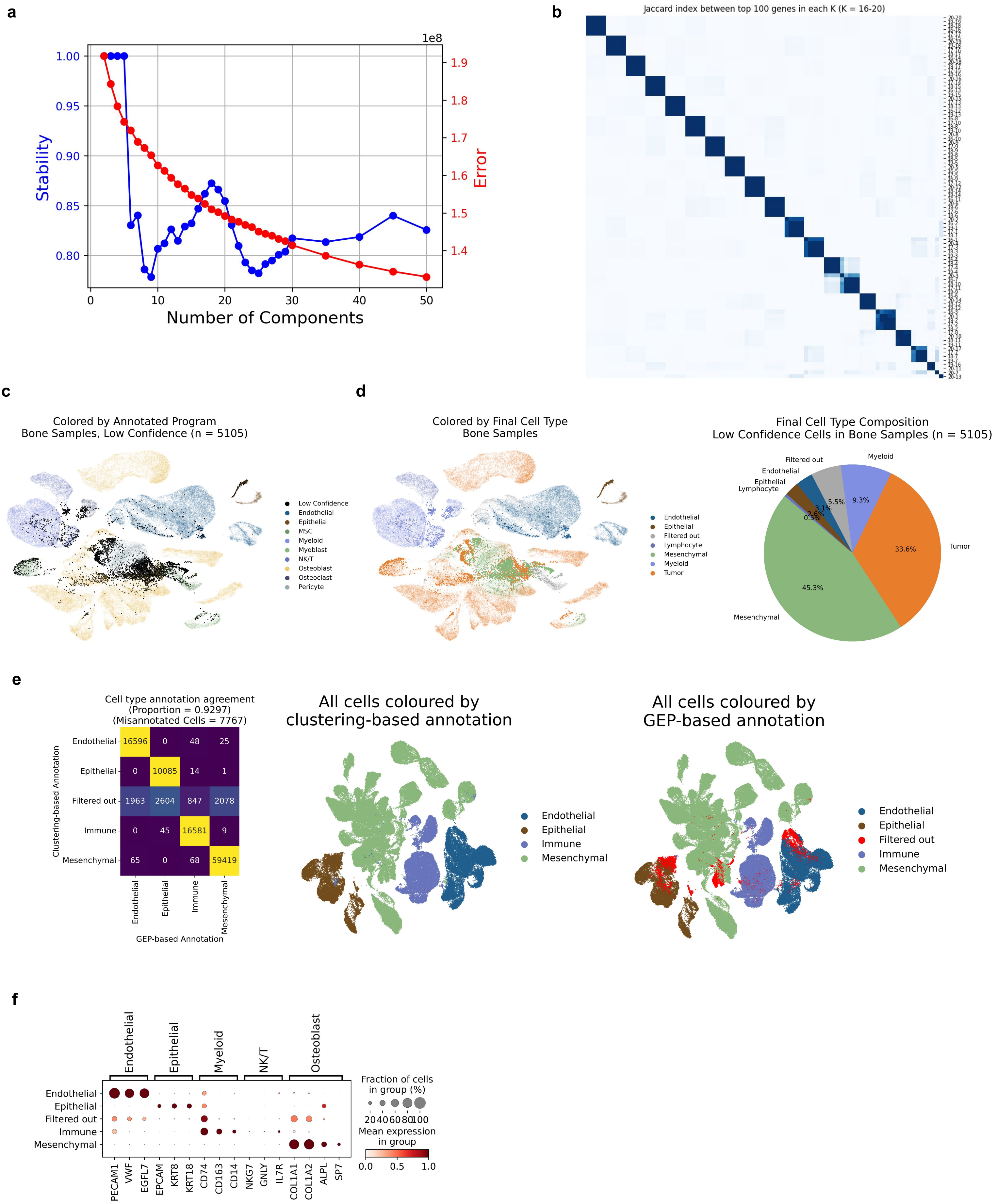

**Supplementary Figure 1: Robustness of cell type annotations to variable parameters**

**(a).** Selection of number of GEPs generated. We select a variable range of gene expression programs (GEPs) (k = 16-20) to generate based on stability and error metrics. **(b).** Evaluation of cNMF stability. The top 100 genes for each program across the variable k runs were computed and clustered according to Jaccard index similarity. **(c).** UMAP visualization of the bone snRNA-seq data, colored by the assigned program. Cells with no dominant cell type program usage was annotated as a "low confidence" cell. **(d).** (Left) UMAP visualization of the bone snRNA-seq data, colored by the final cell type annotation made. (Right) We find that a substantial proportion of the "low confidence" cells were annotated as mesenchyme or malignant. **(e).** (Left) Overlap between annotated cell types of the clustering and GEP-based approaches. (Center) UMAP visualization of the whole cohort colored by clustering-based cell type annotation (Right) UMAP visualization of the whole cohort colored by GEP-based cell type annotation approach. **(f).** Expression of canonical marker genes by annotated cell type. The filtered out cells express multiple cell type markers, suggesting their "doublet"-like cell type.

#### Supplementary Figure 2

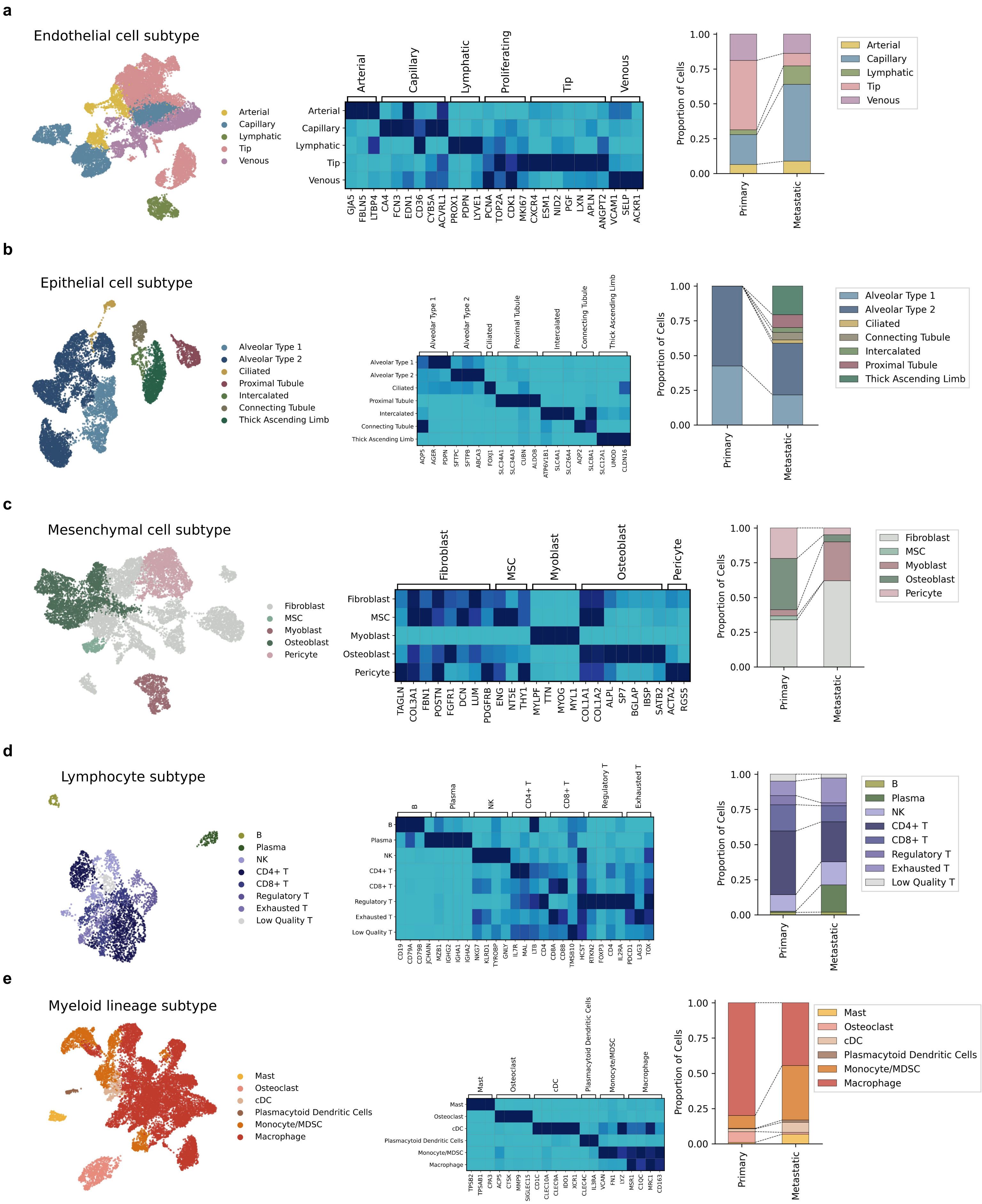

**Supplementary Figure 2: Tumor microenvironment subtype annotation**

UMAP visualization (left), matrix plot of marker gene expression (middle), and proportion plots of subtype cells between the primary / metastatic samples (right) for each of the broad TME cell type classes. Endothelial **(a)**, epithelial **(b)**, mesenchyme **(c)**, lymphocytes **(d)**, and myeloid **(e)**.

### Supplementary Figure 3

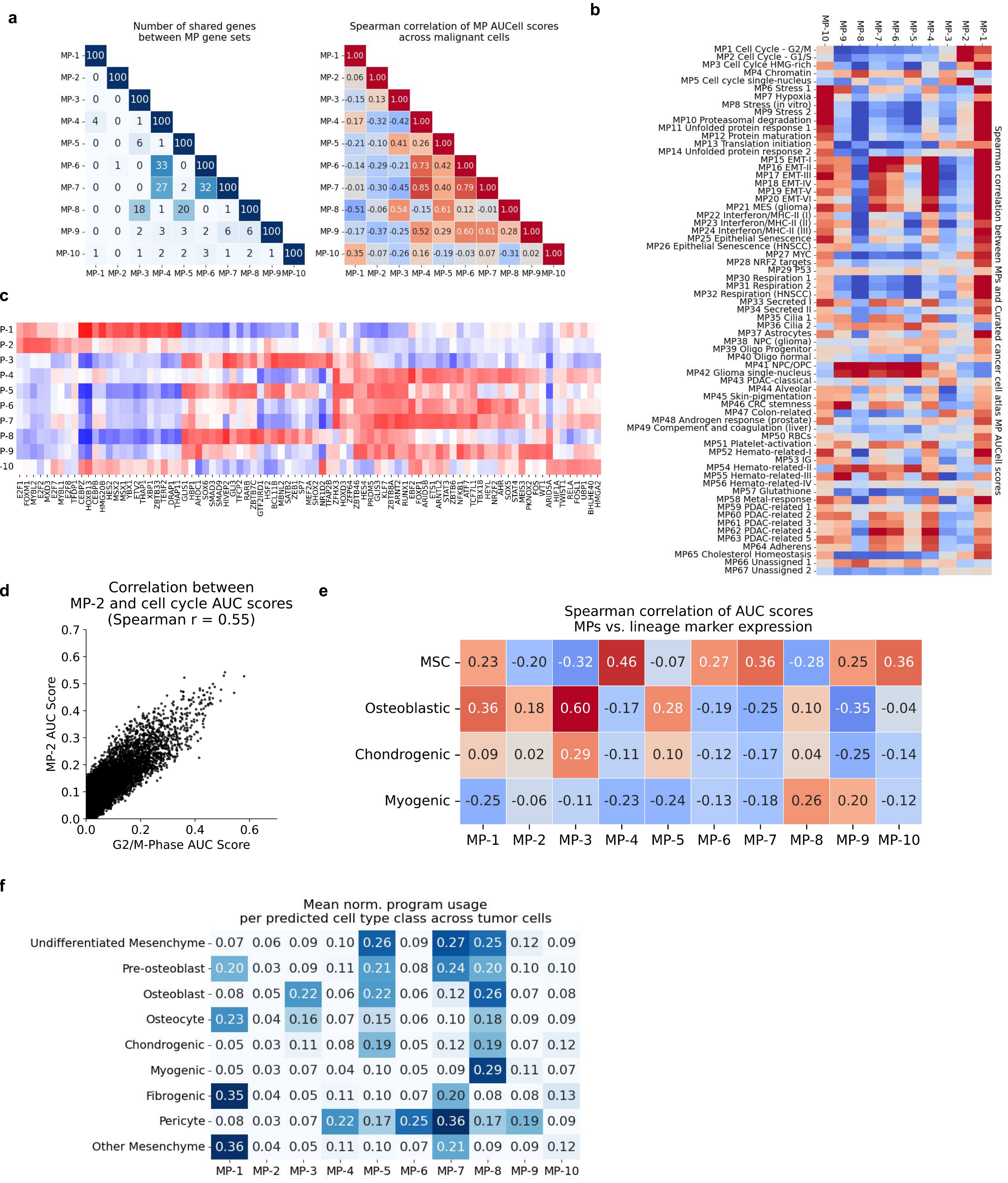

**Supplementary Figure 3: Characterization of all MPs identified across OS malignant cells**

**(a).** Heatmap displaying the pairwise Pearson correlation of meta-program (MP) AUCell scores across all tumor nuclei. **(b).** Correlation matrix aligning the identified MPs with established 3CA meta-programs. **(c).** Heatmap of transcription factor (TF) regulon activity correlated with MP usage. The non-osteoblastic programs MP-5 and MP-8 share a highly active regulatory network, characterized by robust enrichment for FOXP1, MEF2C, PBX3, and ZEB1 regulons. **(d).** Scatter plots depicting the correlation of MP-2 against the G2/M phase cell cycle signature AUCell score. **(e).** Spearman correlation between the MPs and various mesenchymal lineage signature scores. **(f).** Distribution of the usage of all MPs by predicted normal proxy cell type. We first predict the normal proxy cell type for each individual tumor cell, and compute the mean score of each program for the cells predicted as that cell lineage.

### Supplementary Figure 4

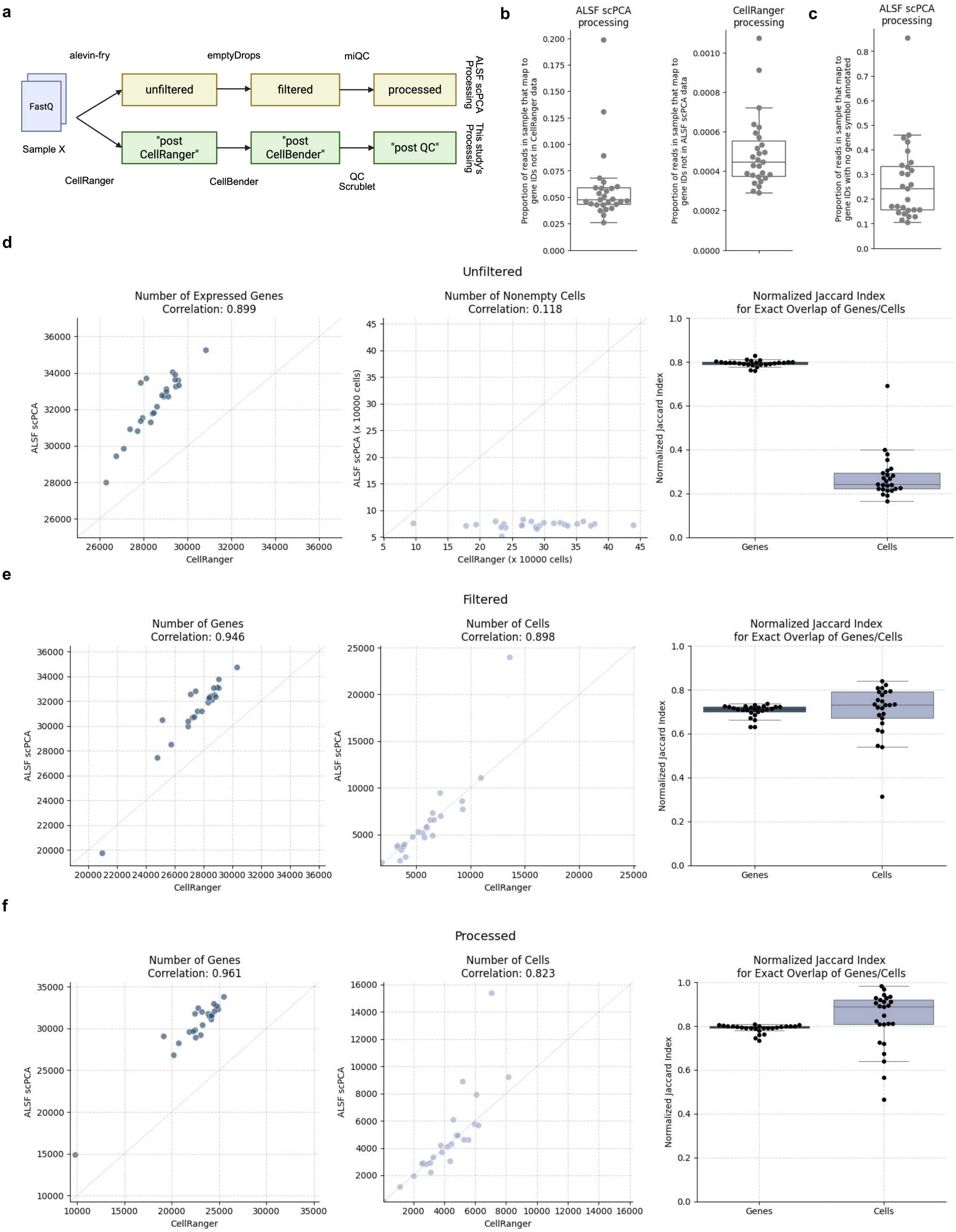

**Supplementary Figure 4: Comparison of this study's processing vs. Alex's Lemonade Stand single-cell Pediatric Cancer Atlas processing of OS snRNA-seq dataset**

**(a).** Schematic overview of the comparable data processing stages between the Alex's Lemonade Stand Foundation (ALSF) scPCA pipeline (alevin-fry) and the pipeline used in this study (CellRanger/CellBender). **(b).** Reference Genome Discrepancies (Absent Genes). (Left) Quantification of reads assigned to genes present in the alevin-fry reference but absent from the CellRanger reference in the alevin-fry processed matrices per sample. (Right) Quantification of reads assigned to genes present in the CellRanger reference but absent from the alevin-fry reference in the CellRanger processed matrices per sample. **(c).** Reference Genome Discrepancies (Unnamed IDs). Comparison of gene IDs between Ensembl 104 (alevin-fry) and Ensembl 98 (CellRanger), highlighting the proportion of reads assigned to unnamed gene IDs in the alevin-fry processed matrices per sample. **(d).** Correlation between scPCA unfiltered counts vs DFCI Post CellRanger counts matrices. (Left) Number of genes expressed in at least one cell per sample (Middle) Number of nonempty cells per sample (Right) Jaccard index for the overlap of genes and cells per sample. **(e).** Correlation between scPCA filtered counts vs DFCI post-CellBender. (Left) Number of genes expressed in at least one cell per sample (Middle) Number of cells passing filtering per sample (Right) Jaccard index for the overlap of genes and cells per sample. **(f).** Correlation between scPCA processed counts vs DFCI post-QC. (Left) Number of genes expressed in at least one cell per sample (Middle) Number of cells passing all QC steps per sample (Right) Jaccard index for the overlap of genes and cells per sample.

Kotliar, D., Veres, A., Nagy, M. A., Tabrizi, S., Hodis, E., Melton, D. A., and Sabeti, P. C. Identifying gene expression programs of cell-type identity and cellular activity with single-cell RNA-seq. 8:e43803. doi: 10.7554/eLife.43803.

Traag, V. A., Waltman, L., and Van Eck, N. J. From louvain to leiden: guaranteeing well-connected communities. 9(1):5233. doi: 10.1038/s41598-019-41695-z.

Wolf, F. A., Angerer, P., and Theis, F. J. SCANPY: large-scale single-cell gene expression data analysis. 19(1):15. doi: 10.1186/s13059-017-1382-0.
